# Joint Learning of Full-structure Noise in Hierarchical Bayesian Regression Models

**DOI:** 10.1101/2021.11.28.470264

**Authors:** Ali Hashemi, Chang Cai, Yijing Gao, Sanjay Ghosh, Klaus-Robert Müller, Srikantan S. Nagarajan, Stefan Haufe

**Author notes:** Ali Hashemi, Klaus-Robert Müller, Srikantan S. Nagarajan, and Stefan Haufe are corresponding authors.

## Abstract

We consider the reconstruction of brain activity from electroencephalography (EEG). This inverse problem can be formulated as a linear regression with independent Gaussian scale mixture priors for both the source and noise components. Crucial factors influencing the accuracy of the source estimation are not only the noise level but also its correlation structure, but existing approaches have not addressed the estimation of noise covariance matrices with full structure. To address this shortcoming, we develop hierarchical Bayesian (type-II maximum likelihood) models for observations with latent variables for source and noise, which are estimated jointly from data. As an extension to classical sparse Bayesian learning (SBL), where across-sensor observations are assumed to be independent and identically distributed, we consider Gaussian noise with full covariance structure. Using the majorization-maximization framework and Riemannian geometry, we derive an efficient algorithm for updating the noise covariance along the manifold of positive definite matrices. We demonstrate that our algorithm has guaranteed and fast convergence and validate it in simulations and with real MEG data. Our results demonstrate that the novel framework significantly improves upon state-of-the-art techniques in the real-world scenario where the noise is indeed non-diagonal and full-structured. Our method has applications in many domains beyond biomagnetic inverse problems.

## I. Introduction

PRECISE knowledge of the noise distribution is a fundamental requirement for obtaining accurate solutions in many regression problems [1], including biomedical imaging applications such as neural encoding models for task-based fMRI analyses [2], [3], electrical impedance tomography (EIT) [4]–[6] or magneto- or electroetoencephalography (M/EEG) inverse problems [7]–[9]. In some of these biomedical imaging applications, however, it is impossible to separately estimate this noise distribution, as distinct “noise-only” (baseline) measurements are not feasible. An alternative is to jointly estimate the regression coefficients and parameters of the noise distribution. This has been pursued both in a (penalized) maximum-likelihood setting (here referred to as *Type-I* approaches) [7] as well as in hierarchical Bayesian settings (referred to as *Type-II*) [8]–[11]. Most contributions in the literature, however, consider only a scalar noise level (homoscedastic noise) or a diagonal noise covariance (i.e., independent between different measurements, heteroscedastic noise) [12]–[14]. These are limiting assumptions in practice as noise may be highly correlated across measurements in many realistic scenarios and, thus, have non-trivial off-diagonal elements.

In this paper, we focus on M/EEG based brain source imaging (BSI), although the proposed algorithm can be used in general regression settings including sparse signal recovery. [15]–[17]. The goal of BSI is to reconstruct brain activity from M/EEG data which can be formulated as a sparse Bayesian learning (SBL) problem. Specifically, we cast it as a linear Bayesian regression model with independent Gaussian scale mixture priors on the parameters and noise. Extending classical SBL approaches, we here consider Gaussian noise with full covariance structure. Prominent sources of correlated noise in M/EEG data are, for example, artifacts caused by eye blinks and the heart beat, muscular artifacts and line noise. Other domains that would benefit from modeling full-structure noise include array processing [18], direction of arrival (DOA) estimation [19], geophysical inverse models [20], and electrical impedance tomography (EIT) [4]–[6].

Algorithms that can accurately estimate noise with full covariance structure in these domains can be expected to achieve more accurate regression models and predictions. This motivates us to present a model and to develop an efficient optimization algorithm for jointly estimating the posterior of regression parameters as well as the noise distribution. More specifically, our contribution in this paper is three-fold:

1. We consider linear regression with Gaussian scale mixture priors on the parameters and *full-structure multivariate Gaussian noise* as opposed to classical SBL approaches that only consider noise distributions with scalar or diagonal structures.
2. We formulate the problem as a hierarchical Bayesian (Type-II maximum-likelihood) regression problem, in which the *source variance hyperparameters* and *a full-structure noise covariance matrix* are *jointly* estimated by maximizing the Bayesian evidence of the model.
3. We derive an efficient algorithm based on the majorization-minimization (MM) framework for jointly estimating the source variances and noise covariance along the Riemannian manifold of positive definite (PD) matrices.

The paper is organized as follows: In Section II, we review the necessary background on Type-II Bayesian learning. We then introduce our proposed algorithm in Section III. Simulation studies and real data analysis demonstrating significant improvement in source localization for EEG/MEG brain source imaging are presented in Sections IV and V, respectively. Finally, Section VI concludes the paper.

## II. Type-II Bayesian Regression

We consider the linear model **Y** = **LX** + **E**, where a set of coefficients or source components, **X**, is mapped to the measurements, **Y**, by forward or design matrix, **L** ∈ ℝ^*M*×*N*^. Depending on the setting, the problem of estimating **X** given **L** and **Y** is called an inverse problem in physics, a multi-task regression problem in machine learning, or a multiple measurement vector (MMV) recovery problem in signal processing [21]. Adopting a signal processing terminology, the *measurement matrix* **Y** ∈ ℝ^*M*×*T*^ captures the activity of *M* sensors at *T* time instants, **y**(*t*) ∈ ℝ^*M*×1^, *t* = 1, …, *T*, while the *source matrix*, **X** ∈ ℝ^*N*×*T*^, consists of the unknown activity of *N* sources at the same time instants, **x**(*t*) ∈ ℝ^*N*×1^, *t* = 1, …, *T*. The matrix **E** = [**e**(1), …, **e**(*T*)] ∈ ℝ^*M*×*T*^ represents *T* time instances of zero-mean Gaussian noise with full covariance **Λ, e**(*t*) ∈ ℝ^*M*×*T*^ ∼ 𝒩 (0, **Λ**), *t* = 1, …, *T*, which is assumed to be independent of the source activations.

The goal of BSI is to infer the underlying brain activity **X** from the EEG/MEG measurement **Y** given a known forward operator, called lead field matrix **L**. In practice, **L** can be computed using discretization methods such as the finite element method (FEM) for a given head geometry and known electrical conductivities [22]. As the number of sensors is typically much smaller than the number of locations of potential brain sources, this inverse problem is highly ill-posed. This problem is addressed by imposing prior distributions on the model parameters and adopting a Bayesian treatment through Maximum-a-Posteriori (MAP) estimation (*Type-I Bayesian learning*) [23]–[27] or, when the model has unknown hyperparameters, through Type-II Maximum-Likelihood estimation (*Type-II Bayesian learning*) [28]–[30]. In this paper, we focus on Type-II Bayesian learning, which assumes a family of prior distributions *p*(**X**|**Θ**) parameterized by a set of hyperparameters **Θ**. These hyper-parameters can be learned from the data along with the model parameters using a hierarchical Bayesian approach [31] through the maximum-likelihood principle:

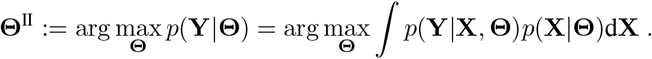

Here we assume a zero-mean Gaussian prior with diagonal covariance **Γ** = diag(***γ***) for the underlying source distribution. That is, **x**(*t*) ∈ ℝ^*N*×1^ ∼ 𝒩 (0, **Γ**), *t* = 1, …, *T*, where ***γ*** = [*γ*_1_, …, *γ*_*N*_] contains *N* distinct unknown variances associated to *N* modeled brain sources. In the Type-II Bayesian learning framework, modeling independent sources through a diagonal covariance matrix leads to sparsity of the resulting source distributions, i.e., at the optimum, many of the estimated source variances are zero. This mechanism is known as *sparse Bayesian learning* (SBL) [31] and is also closely related to the concept of *automatic relevance determination* (ARD) [32] and *kernel Fisher discriminant* (KFD) [33]. Just as most other approaches, SBL makes the simplifying assumption of statistical independence between time samples. This leads to the following expression for the distribution of the sources and measurements:

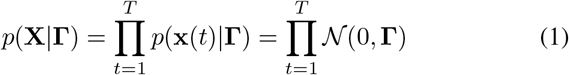

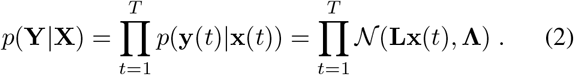

The parameters of the Type-II model are the unknown source variances and the noise covariance, i.e., Θ = {**Γ, Λ**} which are optimized based on the current estimates of the source variances and noise covariance in an alternating iterative process. Given initial estimates of **Γ** and **Λ**, the posterior distribution of the sources is a Gaussian of the form [34]

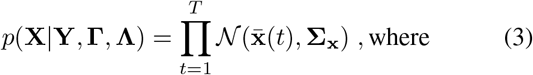

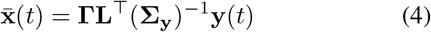

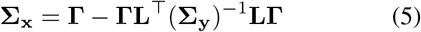

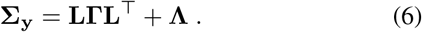

The estimated posterior parameters 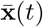 and **Σ**_**x**_ are then in turn used to update **Γ** and **Λ** as the minimizers of the negative (marginal) log-likelihood − log *p*(**Y**|**Γ, Λ**) given by [35]:

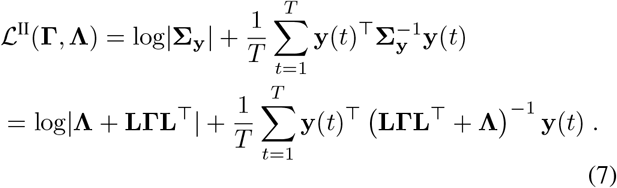

Given the final solution of hyperparameters Θ^II^ = {**Γ**^II^, **Λ**^II^}, the posterior source distribution is obtained by plugging these estimates into (2)–(5).

## III. Proposed Method

### Full-structure Noise (FUN) Learning

Here we propose a novel and efficient algorithm, full-structure noise (*FUN*) learning, which is able to learn the full covariance structure of the noise jointly within the Bayesian Type-II regression framework. We adopt the SBL assumption for the sources, leading to **Γ**-updates previously described in the BSI literature under the name Champagne [28]. As a novelty and main focus of this paper, we here equip the SBL framework with the capability to jointly learn full noise covariances by invoking efficient methods from Riemannian geometry, in particular the geometric mean.

Note that the Type-II cost function in (7) is non-convex and thus non-trivial to optimize. A number of iterative algorithms such as *majorization-minimization* (MM) approaches [36] have been proposed to address this challenge. Following the MM scheme, we here first construct convex surrogate functions that *majorizes* ℒ^II^(**Γ, Λ**) in each iteration of the optimization algorithm. Then, we show the minimization equivalence between the constructed majoring functions and (7). This result is presented in the following theorem:

#### Theorem 1.

*Let* **Λ**^*k*^ *and* 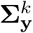 *be fixed values obtained in the* (*k*)*-th iteration of the optimization algorithm minimizing* ℒ^*II*^(**Γ, Λ**). *Then, optimizing the non-convex Type-II ML cost function in* (7), ℒ^*II*^(**Γ, Λ**), *with respect to* **Γ** *is equivalent to optimizing the following convex function, which* majorizes (7):

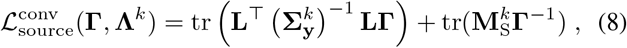

*where* 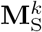 *is defined as:*

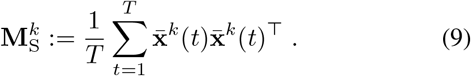

*Similarly, optimizing* ℒ^*II*^(**Γ, Λ**) *with respect to* **Λ** *is equivalent to optimizing the following convex majorizing function:*

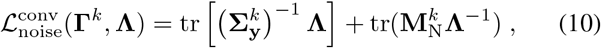

*where* 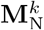 *is defined as:*

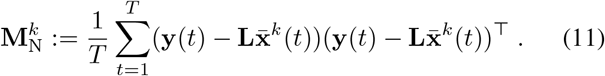

*Proof*. The proof is presented in Appendix A.

We continue by considering the optimization of the cost functions 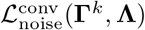 and 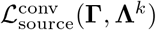 with respect to **Λ** and **Γ**, respectively. Note that in case of noise covariances with full structure, the solution of 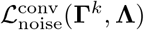 with respect to **Λ** lies within the (*M* ^2^ +*M*)*/*2 Riemannian manifold of PD matrices of size *M*× *M*. This enables us to invoke efficient methods from Riemannian geometry (see [37]), which ensure that the solution at each step of the optimization is contained within the lower-dimensional solution space. Specifically, in order to optimize for the noise covariance, the algorithm calculates the geometric mean between the previously obtained statistical model covariance, 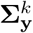, and the empirical sensor-space residuals, 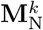, in each iteration. Regarding the solution Of 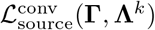, note that we adopt the SBL assumption for the sources by imposing a diagonal structure on the source covariance matrix, **Γ** = diag(***γ***), where ***γ*** = [*γ*_1_, …, *γ*_*N*_]. The update rules obtained from this algorithm are presented in the following theorems:

#### Theorem 2.

*The cost function* 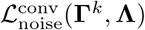 *is strictly geodesically convex with respect to the PD manifold, and its minimum with respect to* **Λ** *can be attained according to the following update rule:*

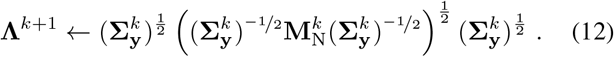

*Proof*. A detailed proof can be found in Appendix B. More-over, a geometric representation of the geodesic path between the pair of matrices 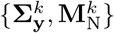 on the PD manifold and the geometric mean between them, representing the update for **Λ**^*k*+1^, is provided in Fig. 1.

**Fig. 1:**
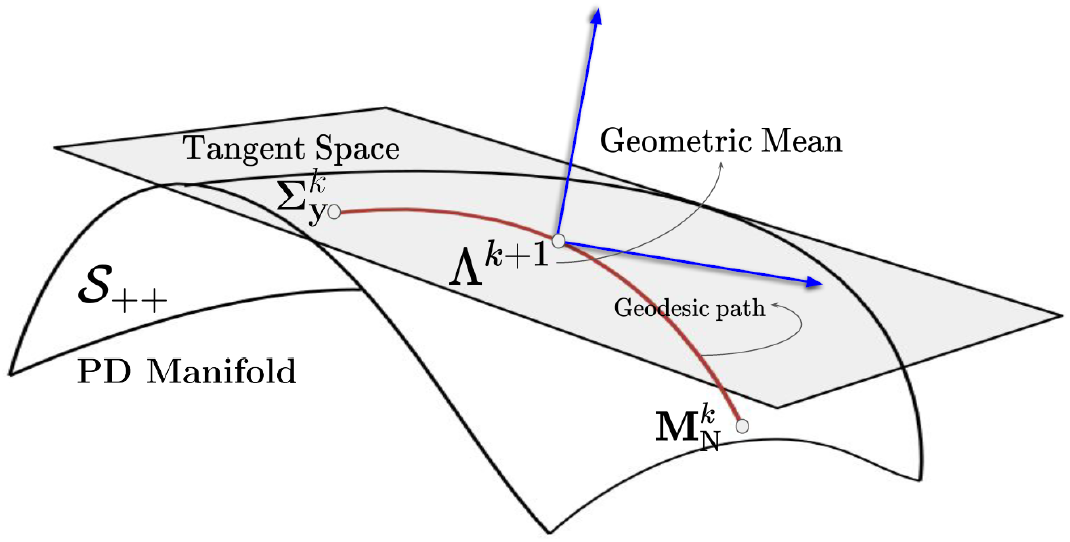
Geometric representation of the geodesic path between the pair of matrices 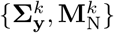 on the PD manifold and the geometric mean between them, which is used to update **Λ**^*k*+1^.

#### Remark 1.

**1**. *Note that the obtained update rule is a closed-form solution for the surrogate cost function*, (10), *which stands in contrast to conventional majorization minimization algorithms (see Section D in the appendix), which require iterative procedures in each step of the optimization*.

#### Theorem 3.

*Constraining* **Γ** *in* (8) *to the set of diagonal matrices with nonnegative elements* 𝒮, *i*.*e*., 𝒮 = {**Γ** | **Γ** = diag(***γ***) = diag([*γ*_1_, …, *γ*_*N*_]^⊤^), *γ*_*n*_ ≥ 0, *for n* = 1,…, *N* },

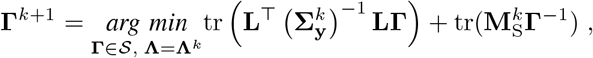

*leads to the following update rule for the source variances:*

**Γ**^*k*+1^ = diag(***γ***^*k*+1^), *where*,

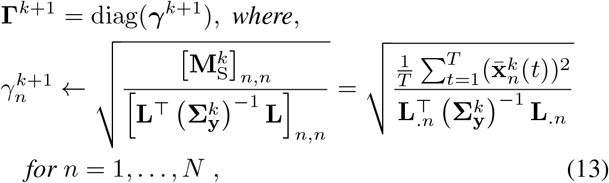

*and where* **L**_.*n*_ *denotes the n-th column of the lead field matrix*.

*Proof*. A detailed proof can be found in Appendix C.

Convergence of the resulting algorithm is shown in the following theorem:

#### Theorem 4.

*Optimizing the non-convex Type-II ML cost function in* (7), ℒ^*II*^(**Γ, Λ**) *with alternating update rules for* **Λ** *and* **Γ** *in* (12) *and* (13) *leads to an MM algorithm with convergence guarantees*.

*Proof*. A detailed proof can be found in Appendix D.

#### Remark 2.

*Note that* (13) *is identical to the update rule of the* Champagne *algorithm [28]. Moreover, various recent Type-II schemes for learning diagonal noise covariance matrices that are rooted in the concept of SBL [8], [9] can also be derived as special cases of FUN learning. Specifically, imposing diagonal structure on the noise covariance matrix for the FUN algorithm, i*.*e*., **Λ** ∈ 𝒮, *results in the noise variance update rules derived in [9] for* heteroscedastic, *and in [8] for* homoscedastic *noise. We explicitly demonstrate the connection between FUN learning and heteroscedastic noise learning in Appendix E*.

#### Remark 3.

*Although FUN is limited to estimating a diagonal source covariance matrix, e*.*g*. **Γ** = diag(***γ***), *this assumption can be relaxed in certain settings. One such setting is when the inverse of* 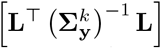 *is well-defined. This is the case whenever the rank of the lead field matrix* **L** *is less than the number of sensors. In the context of BSI, this scenario, for example, occurs when a region-level lead field – instead of a voxel-level lead field – is used. Under this condition, an update rule similar to* (12) *can be obtained for the full-structure source covariance matrix:*

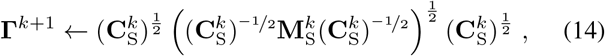

*where* 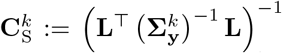. *For additional extensions to other scenarios, please see the discussion section*.

Summarizing, similar to Champagne and other SBL algorithms, the FUN learning approach also assumes independent Gaussian distributed sources with diagonal source covariances, which are updated through (13). As an extension to the classical SBL setting, which assumes the noise distribution to be known, FUN models noise with full covariance structure, which is updated using (12). We summarize the algorithm in Algorithm 1.

#### Remark 4.

*The theoretical results presented in Section III have been obtained for the scalar setting of voxels, where the orientations of the dipolar brain source are assumed to be perpendicular to the surface of the cortex and, hence, only the scalar deflection of each source along the fixed orientation needs to be estimated. In real data, surface normals are hard to estimate or even undefined in case of volumetric reconstructions. Consequently, we model each source here as a full 3-dimensional current vector. This is achieved by introducing three variance parameters for each source within the source covariance matrix*, 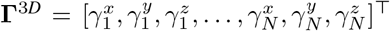. *As all Type-II algorithms considered here model the source covariance matrix* **Γ** *to be diagonal, the proposed extension to 3D sources with free orientation is readily applicable. Correspondingly, a full 3D leadfield matrix*, **L**^3*D*^ ∈ ℝ^*M*×3*N*^,

#### Algorithm 1: Full-structure noise (FUN) learning

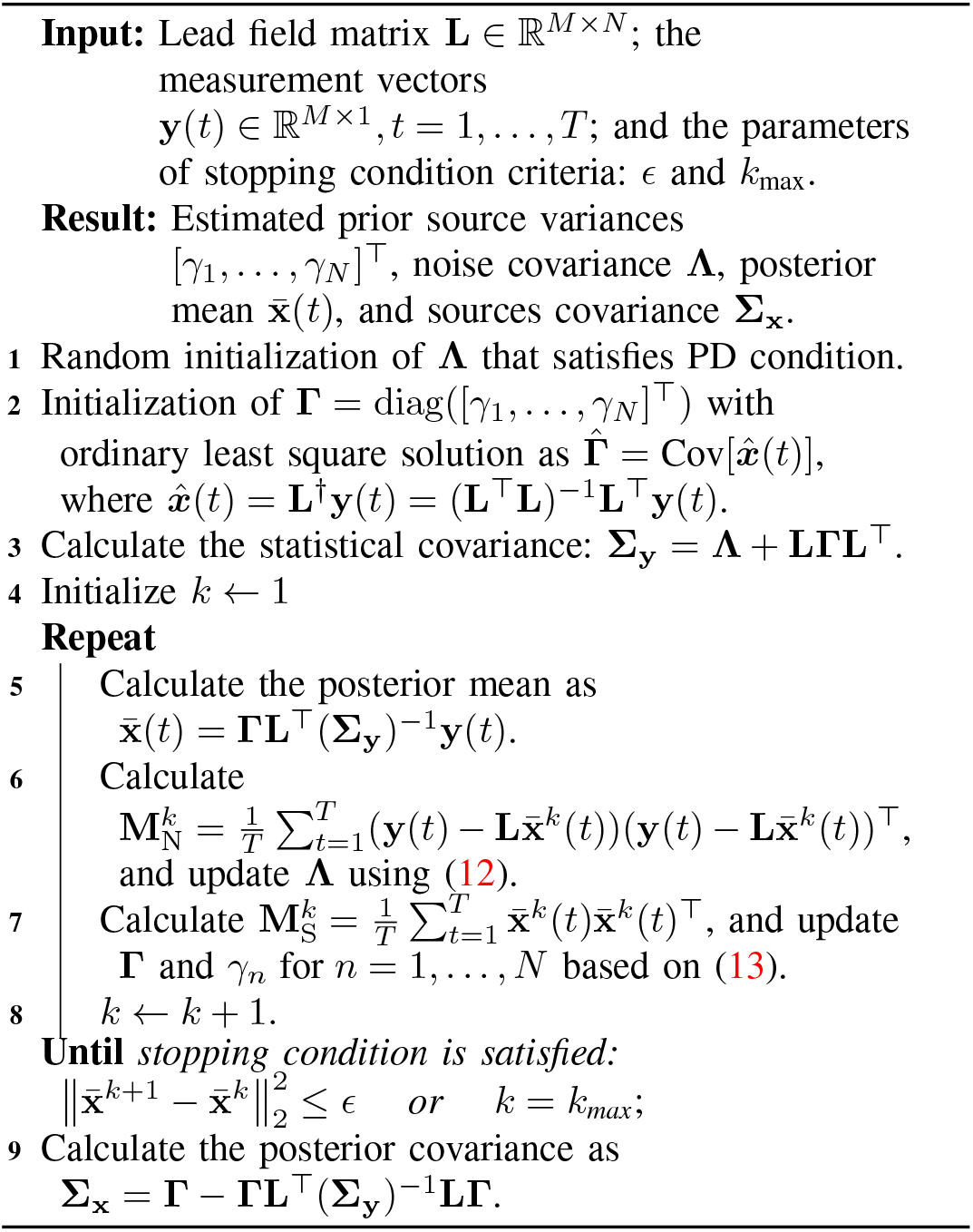

*is used, where we define* **L**^3*D*^ = [**L**_1_, …, **L**_*N*_], *and where N is the number of voxels under consideration and* 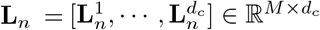 *is the leadfield matrix for n-th voxel with d*_*c*_ *orientations. The k-th column of* **L**_*n*_, *i*.*e*. 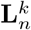 *for k* = 1, …, *d*_*c*_, *represents the signal vector that would be observed at the scalp given a unit current source or dipole at the n-th voxel with a fixed orientation in the k-th direction. The voxel dimension d*_*c*_ *is commonly set to 3 for EEG, and MEG with realistic volume conductor models, and 2 for MEG with single spherical shell models. The update rule in* (13) *can then be reformulated as follows:*

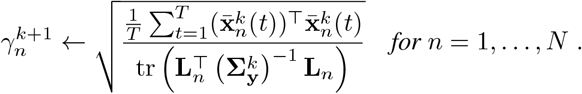

### Complexity Analysis

Suppose FUN takes *K* iterations to converge. The key steps within each iteration of FUN include matrix multiplications of different dimensions, additions of matrices, and a matrix inversion. Of note, **Γ** is diagonal matrix in our setting; which significantly reduces the computational burden. Finally, by retaining only dominating factors and using that *T* ≪ *N, M* ≪ *N*, and log(*N*) *< M* in typical BSI settings, we obtain the overall complexity as 𝒪(*MNT*) + 𝒪(*M* ^2^*N*). Note that since the model used in FUN learning better captures the structure of the noise in most settings, it converges faster than a less accurate diagonal noise model (please see convergence plots in Fig. 2). Therefore, this observation can be interpreted as a trade-off in which even though per-iteration complexity is more significant for FUN compared to heteroscedastic or homoscedastic noise learning variants, fewer iterations are required for FUN to meet the convergence criteria. This behavior can reduce the overall computational complexity of FUN learning and result in a competitive or only slightly increased total computational time compared to diagonal heteroscedastic or homoscedastic noise learning.

**Fig. 2:**
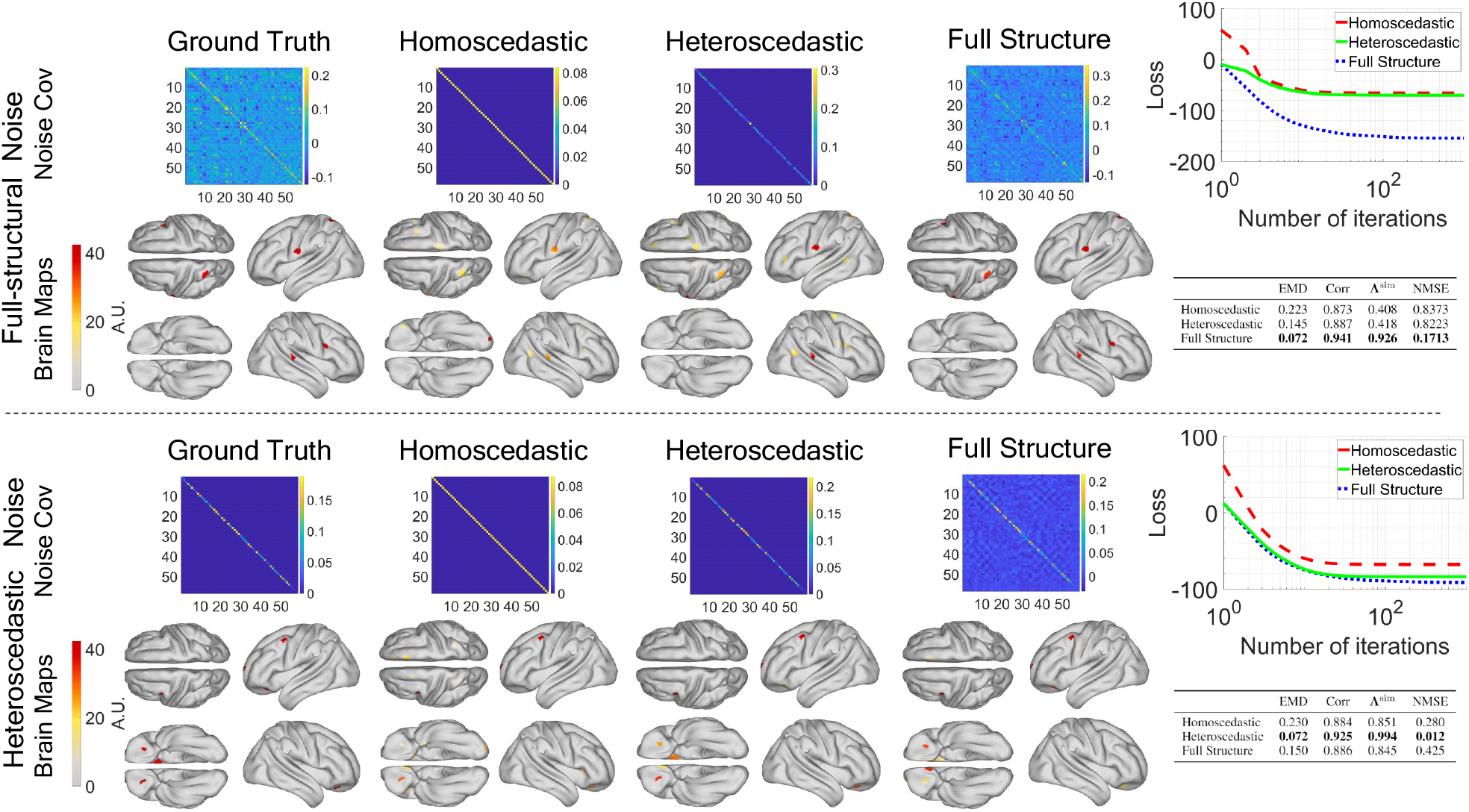
Two examples of the simulated data with five active sources in presence of full-structure noise (upper panel) as well as heteroscedastic noise (lower panel) at 0 dB SNR. Topographic maps depict the locations of the ground-truth active brain sources (first column) along with the source reconstruction results of three noise learning schemes assuming noise with homoscedastic (second column), heteroscedastic (third column), or full structure (fourth column). For each algorithm, the estimated noise covariance matrix is also plotted above the topographic maps. The source reconstruction performance of these examples in terms of EMD and time course correlation (Corr) is summarized in the associated table next to each panel. Beside these two source reconstruction metrics, we also report the accuracy with which the ground-truth noise covariance was estimated in terms of the **Λ**^sim^ and NMSE metrics. The convergence behaviour of all three noise estimation approaches is also shown. Note that the full-structure noise learning approach converges to better minima of the negative log-likelihood than competing approaches regardless of whether the ground-truth noise covariance has full or heteroscedastic structure. However, an advantage in terms of reconstruction is only observed in the former case.

## IV. Numerical Simulations

In this section, we compare the performance of the proposed algorithm to variants employing simpler (home- and heteroscedastic) noise models through an extensive set of simulations. We consider a standard EEG inverse problem, where brain activity is reconstructed from simulated pseudo-EEG data [38]. Our MATLAB codes are publicly accessible at: https://github.com/AliHashemi-ai/FUN-Learning.

### A. Pseudo-EEG Signal Generation

#### Forward Modeling

We used a realistic volume conductor model (of human head) which exhibits a linear relationship between primary electrical source currents generated within the populations of pyramidal neurons in the cortical gray matter [22] and the resulting scalp surface potentials captured by EEG electrodes. The lead field matrix **L** ∈ ℝ^58×2004^ consists of 2004 dipolar current sources and 58 sensors was generated using New York Head model [39]. The orientation of all source currents was fixed to be perpendicular to the cortical surface, so that only scalar source amplitudes needed to be estimated.

#### Source and Noise Model

We simulated a sparse set of *N*_0_ = 5 active sources placed at random locations on the cortex. Neural activity of these sources **X** = [**x**(1), …, **x**(*T*)], *T* = 200 were simulated by sampled from an identically and independently distributed (i.i.d) Gaussian distribution. Gaussian additive noise was randomly sampled from a multivariate zero-mean Gaussian distribution with *full* covariance matrix **Λ**: **e**(*t*) ∈ ℝ^*M*×1^ ∼ 𝒩 (0, **Λ**), *t* = 1, …, *T*. This setting is further referred to as *full-structure noise*. To further investigate the effect of model violation, we generated noise with diagonal covariance matrix, referred as *heteroscedastic noise*. The noise matrix **E** = [**e**(1), …, **e**(*T*)] ∈ ℝ^*M*×*T*^ is normalized and added to the signal matrix **Y**^signal^ = **LX** as follows:

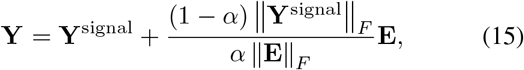

where *α* determines signal-to-noise ratio (SNR) in sensor space defined as SNR = 20log_10_ (*α/*1−*α*). The following SNR (dB) values were used in our experiments: {−12, −7.4, −5.4, −3.5, −1.7, 0, 1.7, 3.5, 5.4, 7.4, 12}.

#### Parameter Initialization

The variances of all voxels were initialized randomly by sampling from a standard normal distribution. The optimization programs were terminated either after reaching convergence (defined by a relative change of the Frobenius-norm of the reconstructed sources between subsequent iterations of less than 10^−8^), or after reaching a maximum of *k*_max_ = 1000 iterations.

#### Performance Metrics

We applied the proposed FUN method on the aforementioned synthetic data to recover the locations and time courses of active brain sources. In addition, two further Type-II Bayesian learning schemes, namely homoscedastic and heteroscedastic Champagne [8], [9], were also included as benchmarks with respect to source reconstruction performance and noise covariance estimation accuracy.

Source reconstruction performance was evaluated according to the following metrics. First, *earth mover’s distance* (EMD) [25], [40], normalized to [0, 1], was used to quantify the spatial localization accuracy. The EMD measures the cost needed to transform two probability distributions defined on the same metric domain (in this case, distributions of the true and estimated sources defined in 3D Euclidean brain space) into each other. Second, the reconstruction error was measured using Pearson correlation between all pairs of simulated and reconstructed (i.e., those with non-zero activations) source time courses. To evaluate the localization error, we also report average Euclidean distance (EUCL) between each simulated source and the best (in terms of absolute correlations) matching reconstructed source.

To assess the recovery of the true support, we computed the F_1_ measure [41]: F_1_ = 2×*TP/*(*P* +*TP* +*FP*), where P denotes the number of true active sources, while TP and FP are the numbers of true and false positive predictions. Note that F_1_ = 1 represents the perfect recovery of the true support.

The performance of the noise covariance estimation was evaluated using tree metrics: Pearson correlation (**Λ**^sim^), the normalized mean squared error (NMSE), which is defined as 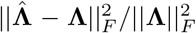, where **Λ** and 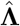 denote true and reconstructed noise covariances, respectively, and finally the log-det Bregman matrix divergence – also known as Stein’s loss – between original and reconstructed noise covariance matrices, denoted by 𝒟_log-det_. An introduction to log-det Bregman matrix divergence in the context of BSI methods can be found in [8, Appendix A]. Note that NMSE measures the reconstruction of the true scale of the noise covariance matrix, while **Λ**^sim^ is scale-invariant and hence only quantifies the overall structural similarity between simulated and estimated noise covariance matrices.

Each simulation was carried out 100 times using different instances of **X** and **E**, and the mean and standard error of the mean (SEM) of each performance measure across repetitions was calculated.

### B. Results

Fig. 2 shows two simulated datasets with five active sources in presence of full-structure noise (upper panel) and heteroscedastic noise (lower panel) at 0 dB SNR. Topographic maps depict the locations of the ground-truth active brain sources (first column) along with the source reconstruction result of three noise learning schemes –noise with homoscedastic, heteroscedastic, and full structure. For each algorithm, the estimated noise covariance matrix is also plotted above the topographic map. Source reconstruction performance was measured in terms of EMD and time course correlation (Corr); and are summarized in the table next to each panel. Besides, the accuracy of the noise covariance matrix reconstruction was measured in terms of **Λ**^sim^ and NMSE.

Fig. 2 (upper panel) allows for a direct comparison of the estimated noise covariance matrices obtained from the three different noise learning schemes. It can be seen that FUN learning can better capture the overall structure of ground truth full-structure noise as evidenced by lower NMSE and similarity errors compared to the heteroscedastic and homoscedastic algorithm variants that are only able to recover a diagonal matrix while enforcing the off-diagonal elements to zero. This results in higher spatial and temporal accuracy (lower EMD and time course error) for FUN learning compared to competing algorithms assuming diagonal noise covariance. This advantage is also visible in the topographic maps.

The lower-panel of Fig. 2 presents analogous results for the setting where the noise covariance is generated according to a heteroscedastic model. Note that the superior spatial and temporal reconstruction performance of the heteroscedastic noise learning algorithm compared to the full-structure scheme is expected here because the simulated ground truth noise is indeed heteroscedastic. The full-structure noise learning approach, however, provides fairly reasonable performance in terms of EMD, time course correlation (corr), and **Λ**^sim^, although it is designed to estimate a full-structure noise covariance matrix. The convergence behaviour of all three noise learning variants is also illustrated in Fig. 2. Note that the full-structure noise learning approach eventually reaches lower negative log-likelihood values in both scenarios, namely full-structure and heteroscedastic noise.

Fig. 3 shows the EMD, the time course reconstruction error, the EUCL and the F1 measure score incurred by three different noise learning approaches assuming homoscedastic (red), heteroscedastic (green) and full-structure (blue) noise covariances for a range of SNR values. The upper panel represents the evaluation metrics for the setting where the noise covariance is full-structure model, while the lower-panel depicts the same metric for simulated noise with heteroscedastic diagonal covariance. Concerning the first setting, FUN learning consistently outperforms its homoscedastic and heteroscedastic counterparts according to all evaluation metrics in particular at low-SNR. Consequently, as the SNR decreases, the gap between FUN learning and the two other variants increases. Conversely, heteroscedastic noise learning shows an improvement over FUN learning according to all evaluation metrics when the simulated noise is indeed heteroscedastic. However, note that the magnitude of this improvement is not as large as observed for the setting where the noise covariance is generated according to a full-structure model and then is estimated using the FUN approach.

**Fig. 3:**
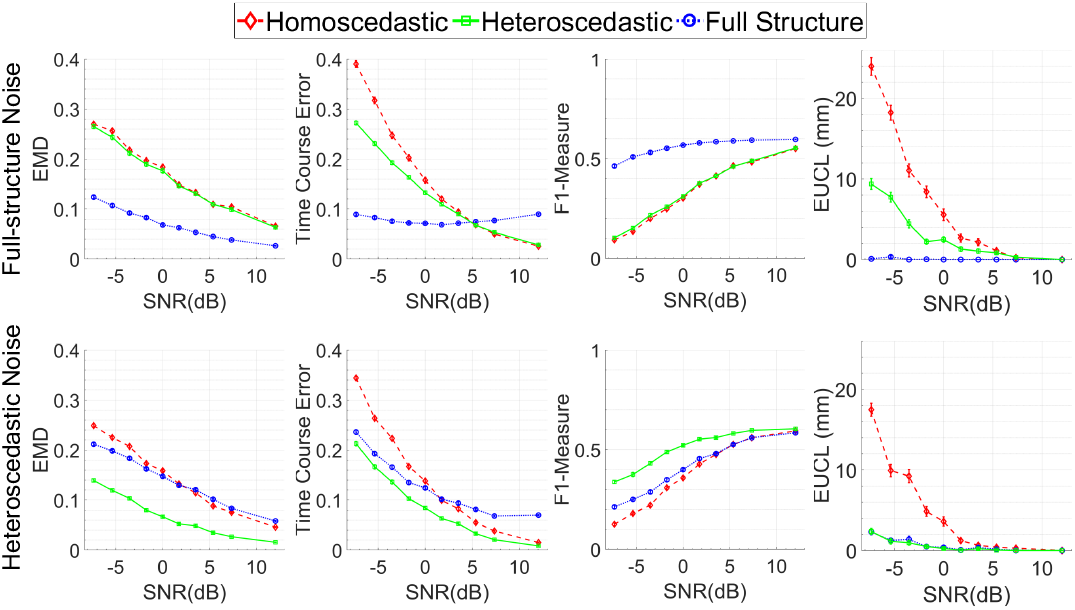
Source reconstruction performance (mean ± SEM) of the three different noise learning schemes for data generated by a realistic lead field matrix. Generated sensor signals were superimposed by either full-structure or heteroscedastic noise covering a wide range of SNRs. Performance was measured in terms of the earth mover’s distance (EMD), time-course correlation error, F1-measure and Euclidean distance (EUCL) in (mm) between each simulated source and the reconstructed source with highest maximum absolute correlation.

Fig. 4 depicts the accuracy if the estimated noise covariance matrix reconstructed by three different noise learning approaches assuming noise with homoscedastic (red), heteroscedastic (green) and full (blue) structure. The ground truth noise covariance matrix either had full (upper row) or heteroscedastic (lower row) structure. Performance was measured in terms of similarity, NMSE, and 𝒟_log-det_. To be consistent with NMSE, we report “similarity error”, defined as 1 − **Λ**^sim^, instead of similarity, **Λ**^sim^. Similar to the trend observed in Fig. 3, full-structure noise learning leads to better noise covariance estimation accuracy (lower NMSE and similarity error) for the full-structure noise model, while superior reconstruction performance is achieved for heteroscedastic noise learning when true noise covariance is heteroscedastic.

**Fig. 4:**
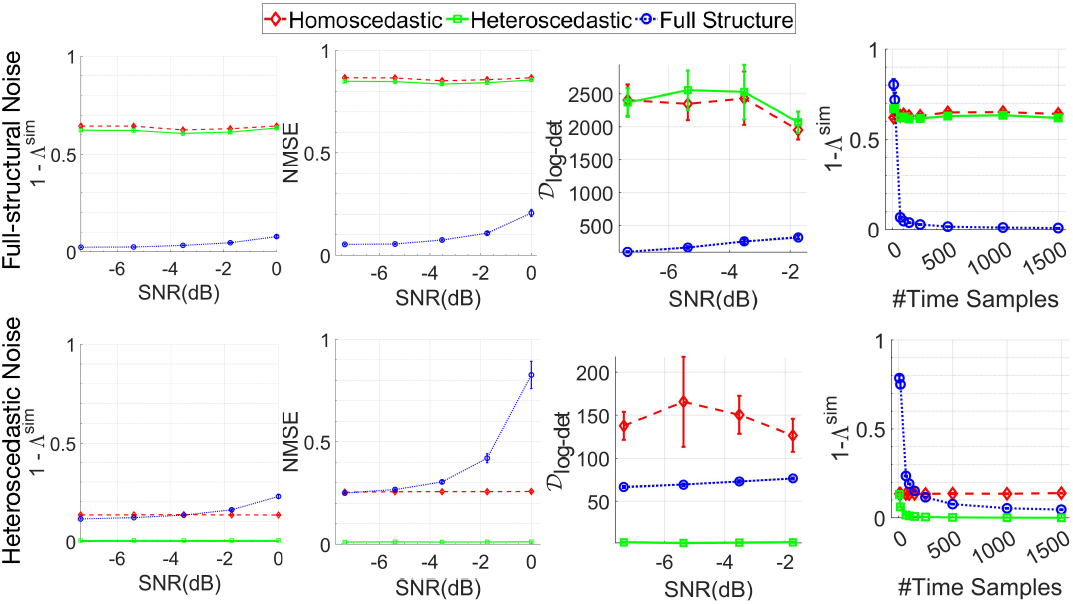
Accuracy of the noise covariance matrix reconstruction incurred by three different noise learning approaches assuming homoscedastic (red), heteroscedastic (green) and full-structure (blue) noise covariances. The ground-truth noise covariance matrix is either full-structure (upper row) or heteroscedastic diagonal (lower row). Performance was assessed in terms of the Pearson correlation between the entries of the original and reconstructed noise covariance matrices, **Λ** and 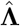, denoted by **Λ**^sim^ (first column). Shown is the similarity error 1 − **Λ**^sim^. Further, the normalized mean squared error (NMSE) between **Λ** and 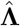, defined as NMSE = 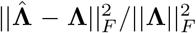 and the log-det Bregman matrix divergence between original and reconstructed noise covariance matrices, denoted by 𝒟_log-det_ are reported (second and third column). The last column depicts the performance of FUN learning as well as heteroscedastic and homoscedastic noise learning for different numbers of time samples as measured by Pearson correlation error between true and reconstructed noise covariance matrices.

The last column of Fig. 4 depicts the performance of FUN learning as well as heteroscedastic and homoscedastic noise learning approaches in terms of the Pearson correlation error, 1 − **Λ**^sim^, for different numbers of time samples. For this experiment, the SNR is set to −3.5 dB and the following number of time samples are used: *T* = {10, 20, 50, 70, 100, 150, 250, 500, 1000, 1500}. The rest of the parameters are set to the values explained in Section. IV-A.

Note that all model inference relies on the robustness of the estimated sample covariance matrix. According to the observed results, we, therefore, conclude that, when the number of samples is small, the sample covariance estimate becomes unreliable and correspondingly will negatively impact the performance of all algorithms. The inference quality, however, can be significantly improved for FUN learning by increasing the number of time samples.

#### Remark 5.

*When the true noise is heteroscedastic, the inference algorithm with a generative model that matches the true scenario, in this case, heteroscedastic noise learning, outperforms a more complex full-structure noise learning model that has many more parameters to be estimated to become zero, i*.*e. more degrees of freedom (DoF). Thus, the full-structure noise (FUN) learning model requires more data to converge to the true model. This behavior is confirmed in the last column of Fig. 4, where we observe that both models, namely heteroscedastic noise learning and FUN, converge at large data lengths when the true noise is heteroscedastic. Notably, while the FUN and heteroscedastic noise learning solutions converge when the true noise is heteroscedastic, the same is not true when the true noise has full-structure. As only FUN learning is able to deal with full structure, its performance is dramatically better than that of heteroscedastic (and homoscedastic) noise learning across all sample sizes in this setting*.

## V. Analysis of Real MEG Data

### A. Auditory and Visual Evoked Fields (AEF and VEF)

All MEG data used here were acquired in the Biomagnetic Imaging Laboratory at the University of California San Francisco (UCSF) with an Omega 2000 whole-head MEG system from CTF Inc. (Coquitlam, BC, Canada) at a sampling rate of 1200 Hz. All human participants provided informed written consent prior to study participation and received monetary compensation for their participation. The studies were approved by the University of California, San Francisco Committee on Human Research.

Lead-fields for each subject were calculated using NUT-MEG [42] assuming a single spherical shell volume conductor model resulting in only two spherical orientations. Lead-fields were constructed at a voxel resolution of 8 mm. Furthermore, each lead-field column was normalized. Neural responses to auditory evoked fields (AEF) and visual evoked fields (VEF) stimulus were localized using the FUN algorithm and other benchmarks. The AEF response was elicited during passive listening to binaural tones (600 ms duration, carrier frequency of 1 kHz, 40 dB SL). The VEF response was elicited while subjects were viewing pictures of objects projected onto a screen and subjects were instructed to overtly name the objects [43], [44]. Up to 120 AEF and 100 VEF trials were collected. For both AEF and VEF data, trials with clear artifacts or visible noise in the MEG sensors that exceeded 10 pT fluctuations were excluded prior to source localization analysis.

Both AEF and VEF data were digitally filtered to a pass-band of 1 to 70 Hz to remove artifacts and DC offset, and time-aligned to the stimulus onset. Averaging was then performed across sets of trials of increasing size: {10, 20, 40, 60, 100} trials for AEF, and {10, 20, 40} trials for VEF analyses. The pre-stimulus window was selected to be 100 ms prior to stimulus onset. The post-stimulus time window for AEF was selected to be +50 ms to +150 ms. For VEF data, we focused on source reconstruction in two time-windows – an early window ranging from +100 ms to +150 ms around the traditional M100 response, and a later time window ranging from +150 ms to +225 ms around the traditional M170 responses [35], [45]–[47].

Fig. 5 shows the reconstruction of the AEF for different number of trial averages for a representative subject using FUN learning along with Type-I and Type-II BSI benchmark methods. In addition to heteroscedastic Champagne, two classical non-SBL source reconstruction schemes were included for comparison. The minimum-current estimate (MCE) algorithm [48] shown here is an example of a sparse Type-I method based on *ℓ*_1_-norm minimization. Additionally, eLORETA [49], represents a smooth inverse solution based on 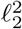-norm minimization.

**Fig. 5:**
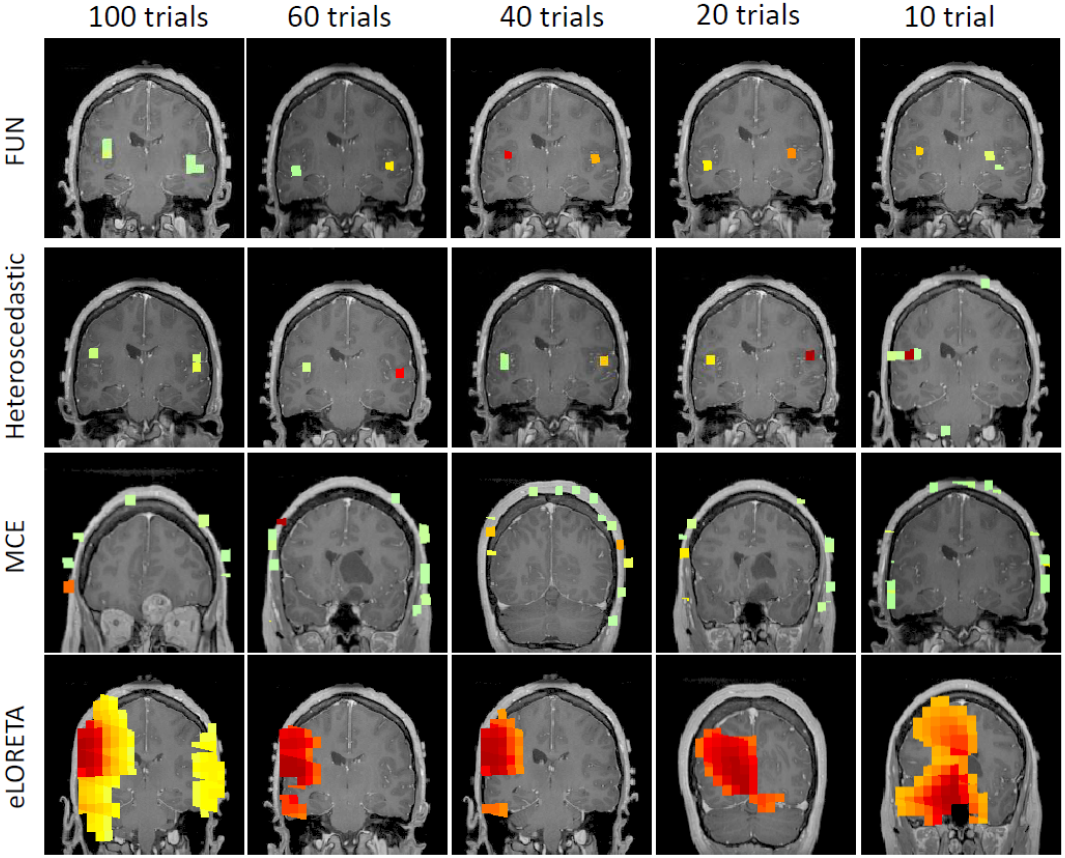
Auditory evoked field (AEF) localization results from one representative subject for different numbers of trial averages using FUN learning, heteroscedastic Champagne, MCE and eLORETA. All reconstructions of FUN learning algorithm show focal sources at the expected locations of the auditory cortex. Even when limiting the number of trials to as few as 10 reconstruction result of FUN learning is accurate, it severely affects the reconstruction performance of competing benchmark methods.

Reconstruction performance of all algorithms for different trial averaging with 10, 20, 40, 60, and 100 trials are shown. All trials were selected randomly prior to averaging. As the subplots for different numbers of trial averages demonstrate, FUN learning can accurately localize bilateral auditory activity to Heschel’s gyrus, the characteristic location of the primary auditory cortex, even with as few as 10 trials. In this challenging setting, FUN outperforms all competing methods.

Regarding the comparison between FUN and the heteroscedastic noise learning approach on real data as demonstrated in Fig. 5, it is not straightforward to evaluate the performance of BSI approaches quantitatively due to the absence of the ground truth. Therefore, the quality of the reconstructions is commonly assessed based on prior neurophysiological knowledge. In Fig. 5, we observed an involvement of both bilateral Heschl’s gyri, which is expected for localization of auditory cortex. Indeed, qualitatively, FUN is able to localize both bilateral auditory activities even when the number of trials is limited to 10. For this setting, the heteroscedastic noise learning approach was only able to locate the left Heschl’s gyrus auditory activity. These results highlight the importance of accurate noise covariance estimation on the fidelity of source reconstructions.

Fig. 6 shows the localization and time series reconstruction of VEF activity for a single subject using FUN and heteroscedastic noise learning Champagne, eLORETA and MCE. Reconstruction performance is again shown for the number of trials used for averaging ranging from 10 to 40. Trials were randomly chosen from the full dataset without replacement prior to averaging. Within each panel, the top shows the source localization of the M100 (1^st^ peak) and M170 (2^nd^ peak) responses, respectively. The time course of the most prominent source (indicated by the intersecting green lines) across a +25 ms to +275 ms window is presented below the source localization results. Blue lines represent the voxel power with arbitrary units averaged across ten independent experiments (that is, ten random selections of trials for trial averaging). Blue shades represent the standard error of the mean (SEM) across different trial averaging experiments. We also included three additional benchmark algorithms, sLORETA [50], S-FLEX [25] and the LCMV beamformer [51] in Fig. 7. In comparison to MCE and eLORETA, FUN shows accurate localization capability, while the former benchmarks did not yield reliable results for averages of only ten trials. Even when the number of trials used for averaging was increased to 20, these benchmarks yielded neither good spatial localization of the two visual cortical peaks, nor were the expected time courses of activation reconstructed. Furthermore, FUN detects two salient and clear peaks in each time window in contrast to other benchmarks, where the salience of the early and late peaks are less prominent. Results obtained from FUN are also robust across different SNRs/numbers of trial averages. For more benchmark results, please see Fig. 7.

**Fig. 6:**
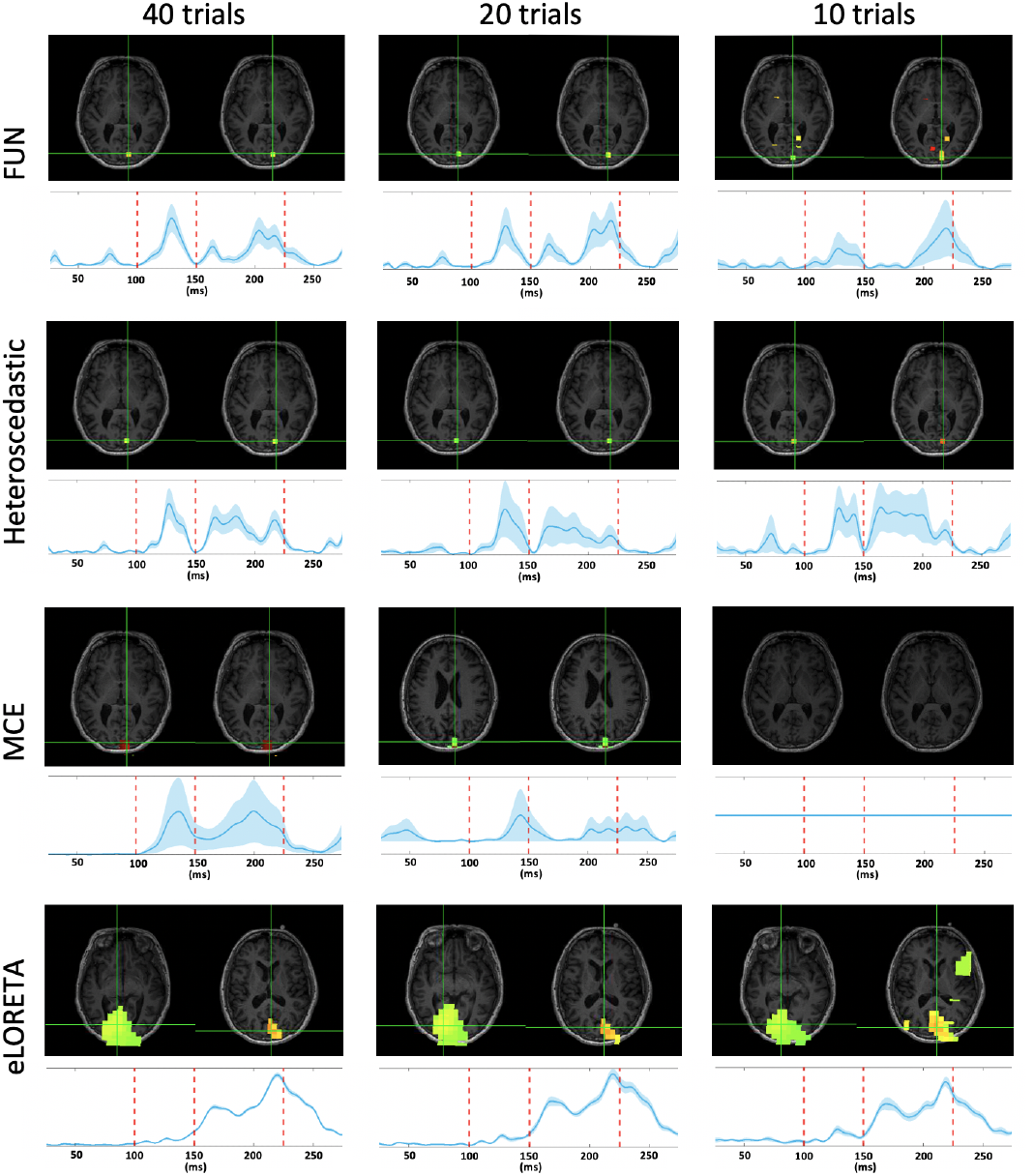
Localization and time series results of visual evoked field (VEF) activity for a single subject using FUN and benchmarks. Comparing with MCE and eLORETA, FUN shows accurate localization capability. Furthermore, FUN detects sharper 2^nd^ peaks when compared to the heteroscedastic noise-learning Champagne, which is consistent with the sharp response of the VEF. The results obtained by FUN are robust across different SNRs/numbers of trial averages. For additional benchmark results, please see Fig. 7.

**Fig. 7:**
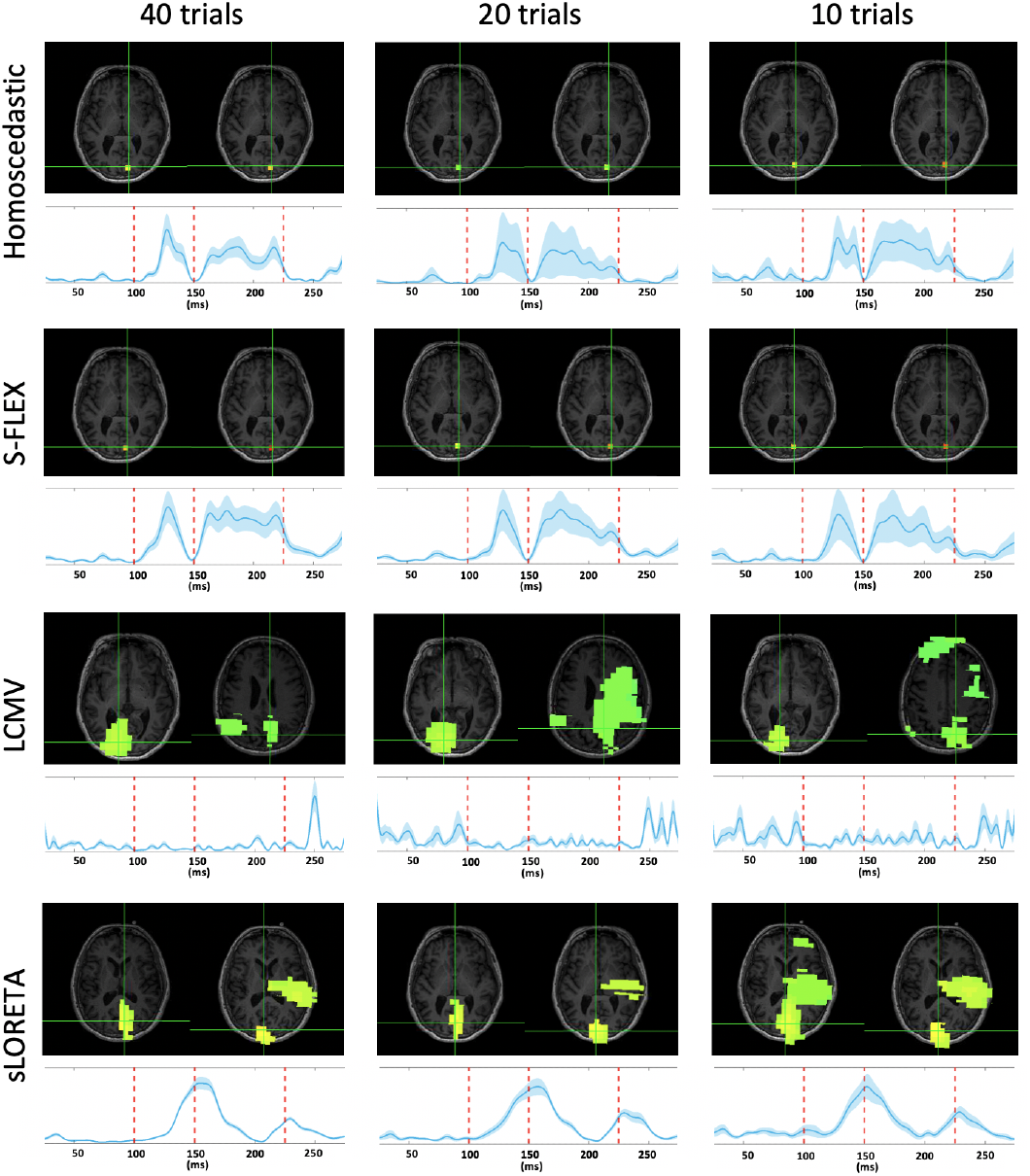
Localization and reconstructed time series of visual evoked field (VEF) activity for a single subject using another four benchmark algorithms. FUN outperforms LCMV beamformer and sLORETA in terms of localization. Moreover, the activation time courses derived from homoscedastic noise learning Champagne and S-FLEX do not exhibit as sharp responses as observed for FUN. The noise level used for S-FLEX reconstructions was set to values learnt from classical Champagne algorithm with noise learning.

### B. Resting-state data

Resting-state data are particularly suited for the FUN algorithm because of the lack of baseline data on which the noise distribution could be estimated. Here, we show that FUN is able to learn the underlying noise distribution and consistently recover brain activity. For this analysis, three subjects were instructed simply to keep their eyes closed and remain awake. We collected four trials per subject, where each trial was one minute long. We randomly chose 30 seconds or equivalently 36000 time samples for brain source reconstruction from one trial of each subject. These resting-state MEG data were digitally filtered using a pass-band ranging from 8 to 12 Hz (alpha band) to remove artifacts and DC offset.

Localization of resting state alpha band activity from the three subjects are shown in Fig. 8. The first three columns show the estimated source covariance patterns (with the application of a threshold of 10% the peak value) for the three noise learning variants of Champagne. Each row represents one subject. The corresponding loss function values across 1000 iterations are shown in the last column. FUN consistently localizes all subjects’ brain activity predominantly near the midline occipital lobe or posterior cingulate gyrus consistent with expected locations of alpha generators known to dominate resting-state activity.

**Fig. 8:**
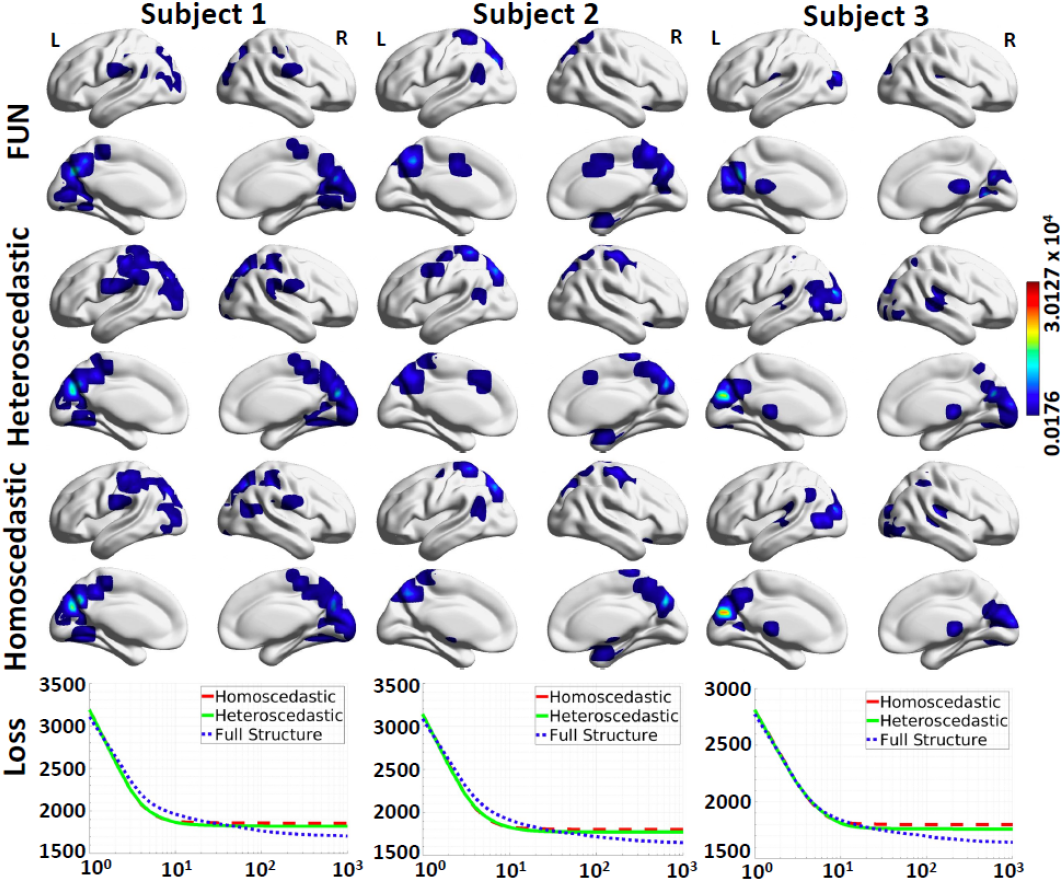
Localization of resting-state brain activity for three subjects using FUN and the heteroscedastic and homoscedastice noise learning variants of Champagne. The source variance patterns estimated by each algorithm are projected onto the cortical surface. The convergence behaviour of all three noise estimation approaches is also shown in terms of the negative log-likelihood cost function. FUN converges to better minima when compared to these benchmarks.

## VI. Discussion

In this paper, we focused on sparse regression within the hierarchical Bayesian regression framework and its application in EEG/MEG brain source imaging. We proposed an efficient optimization algorithm for jointly estimating Gaussian regression parameter distributions as well as Gaussian noise distributions with full covariance structure within a hierarchical Bayesian framework. Using the Riemannian geometry of positive definite matrices, we derived an efficient algorithm for jointly estimating brain source variances and noise covariance. The benefits of our proposed framework were evaluated within an extensive set of experiments in the context of the electromagnetic brain source imaging inverse problem and showed significant improvement upon state-of-the-art techniques in the realistic scenario where the noise has full covariance structure. The practical performance of our method is further assessed through analyses of real auditory evoked fields (AEF), visual evoked fields (VEF) and resting-state MEG data.

In the context of BSI, [52] proposed a method for selecting a single regularization parameter based on cross-validation and maximum-likelihood estimation, while [53]–[57] assume more complex spatio-temporal noise covariance structures. A common limitation of these works is, however, that the noise level is not estimated as part of the source reconstruction problem on task-related data but from separate noise recordings. Our proposed algorithm substantially differs in this respect, as it learns the noise covariance jointly with the brain source distribution from the same data. This joint estimation perspective is opposed to a step-wise independent estimation process that can cause to error accumulation. The idea of joint estimation of brain source activity and noise covariance has been previously proposed for Type-I learning methods in [7], [58]. Bertrand et al. [7] proposed a method to extend the group Lasso class of algorithms to multi-task learning, where the noise covariance is estimated using an eigenvalue fit to the empirical sensor space residuals defined as 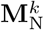 in Theorem 1. In contrast, FUN learning uses Riemannian geometry principles, e.g., the geometric mean between the sensor space residuals 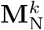 and the previously obtained statistical model covariance, 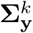. This enables us to robustly estimate the noise covariance as part of the model, in contrast to the method proposed in [7], which estimates the noise covariance solely based on the eigenvalues of the observed sensor space residuals. Furthermore, in contrast to these Type-I likelihood estimation methods, FUN is a Type-II method, which learns the prior source distribution as part of the model fitting. Type-II methods have been reported to yield results that are consistently superior to those of Type-I methods [8], [9], [46], [47], [59]. Our numerical results show that the same holds also for FUN learning, which performs on par or better than existing variants from the Type-II family (including conventional Champagne) in this study.

The question of which noise model to use on real data can be addressed through well-known model selection techniques from the machine learning literature. One such strategy is to evaluate the Type-II negative log-likelihood loss of both models and pick the model that achieves the lowest loss, i.e. choose models that maximize the Bayesian evidence. This was the objective of our analysis in Fig. 8, where we demonstrated that the localization of resting-state brain activity using FUN learning converges to a lower negative log-likelihood loss, i.e., better Bayesian model evidence, than heteroscedastic noise learning, which indicates the superiority of FUN learning and the necessity to model full-structure noise. Furthermore, it is also possible to evaluate the Type-II likelihood, or, the Bayesian model evidence, out-of-sample in order to perform model selection in real data analyses. This approach may be suitable when parameters of the Type-II likelihood are being optimized as is the case here for all approaches. Using this technique, which was successfully employed in [8], the data samples are first split into two parts, namely the training set and the testing set, i.e., hold-out data. For real data analysis, the data can be split among different trials or sensor subsets. The model parameters are fitted to the training set and the Type-II (Bregman) or Type-I likelihoods of the fitted model are then evaluated on the hold-out data (see [8, Eqs. 31 and 32] for related formulations). Note that since the hold-out data are not used during model fitting, the likelihood evaluation on this data is called *out-of-sample* likelihood. The BSI method that achieves better out-of-sample likelihood with respect to the evaluation metric can be considered superior in terms of performance for real data analysis. Formal comparisons of the performance of these model selection techniques on different real data sets are interesting explorations and are considered one of the directions of our future work.

Noise learning has also attracted attention in functional magnetic resonance imaging (fMRI) [2], [3], [60], where various models like matrix-normal (MN), factor analysis (FA), and Gaussian-process (GP) regression have been proposed. The majority of the noise learning algorithms in the fMRI literature rely on the EM framework, which is quite slow in practice [8] and has convergence guarantees only under certain restrictive conditions [36], [61]–[63]. In contrast to these existing approaches, our proposed framework not only applies to the models considered in these papers, but also benefits from theoretically proven convergence guarantees. To be more specific, we showed in this paper that FUN learning is an instance of the wider class of majorization-minimization (MM) framework, for which provable fast convergence is guaranteed. It is worth emphasizing our contribution within the MM optimization context as well. Unlike many other MM implementations, where surrogate functions are minimized using an iterative approach, our proposed algorithm is more efficient because it obtains a closed-form solution for the minimum of the surrogate function in each step.

As pointed out in the introduction, electrical impedance tomography (EIT) is another practical example in which the noise interference is highly correlated across measurements; and thus, indeed has full covariance structure. The authors in [4], [5] addressed this problem using SBL techniques for multiple measurement vector (MMV) models. Since the noise in these works is restricted to scalar or diagonal covariance structure, FUN learning could be used to model more realistic full-structural noise also in EIT problems.

While being broadly applicable (see [64, Appendix A] for a comprehensive list of potential applications), our approach is nevertheless limited by a number of factors. Although Gaussian noise distributions are commonly justified, it would be interesting to include more robust non-Gaussian noise distributions in our framework. Besides, signals in real-world scenarios often lie in a lower-dimensional space compared to the original high-dimensional ambient space due to the correlations that exist in the data. Therefore, imposing physiologically plausible constraints on the noise model, e.g., low-rank, Toeplitz, or Kronecker structure [65], [66], not only provides side information that can be leveraged for the reconstruction but also reduces the computational cost in two ways: a) by reducing the number of parameters and b) by taking advantage of efficient implementations using circular embeddings and the fast Fourier transform [67], [68]. In our recent work [68], we employed separable Gaussian distributions using Kronecker products of temporal and spatial covariance matrices. The proposed efficient algorithms exploit the intrinsic Riemannian geometry of temporal autocovariance matrices. For stationary dynamics described by Toeplitz matrices, the theory of circulant embeddings was employed.

### A. Proof of Theorem 1

*Proof*. We start the proof by recalling (7):

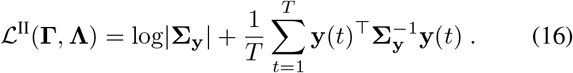

The upper bound on the log |**Σ**_**y**_| term can be directly inferred from the concavity of the log-determinant function and its firstorder Taylor expansion around the value from the previous iteration, 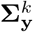, which provides the following inequality [36, Example 2]:

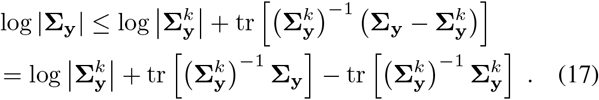

Note that the first and last terms in (17) do not depend on **Γ**; hence, they can be ignored in the optimization procedure. Now, we decompose **Σ**_**y**_ into two terms, each of which only contains either the noise or source covariances:

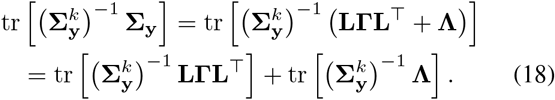

In next step, we decompose the second term in (7), 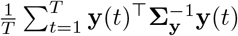, into two terms, each of which is a function of either only the noise or only the source covariances. To this end, we exploit the following relationship between sensor and source space covariances:

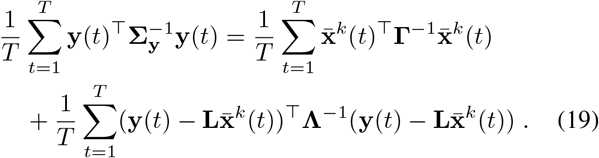

By combining (18) and (19), rearranging the terms, and ignoring all terms that do not depend on **Γ**, we have:

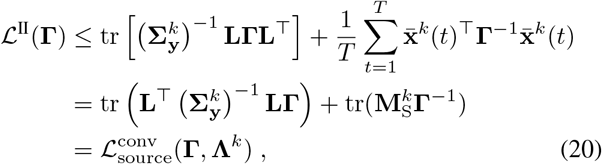

where 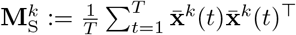.

This proves the equivalence of (7) and (8) when the optimization is performed with respect to **Γ**.

The equivalence of (7) and (10) can be shown analogously, with the difference that we only focus on noise-related terms in (18) and (19):

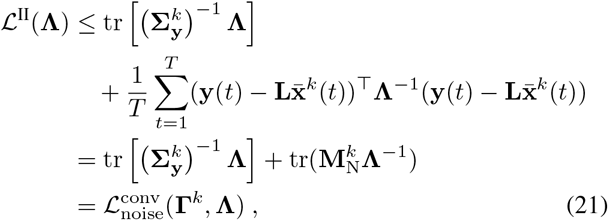

where 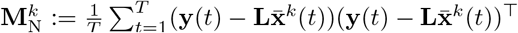.

Summarizing, we have shown that optimizing (7) is equivalent to optimizing 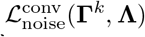 and 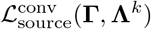, which concludes the proof. □

### B. Proof of Theorem 2

Before presenting the proof, the subsequent definitions and propositions are required:

#### Definition 1

(Geodesic on the positive definite (PD) manifold). *Let be a Riemannian manifold, i*.*e*., *a differentiable manifold whose tangent space is endowed with an inner product that defines local Euclidean structure. Then, a geodesic between two points on* ℳ, *denoted by* **p**_0_, **p**_1_ ∈ ℳ, *is defined as the shortest connecting path between those two points along the manifold, ζ*_*l*_(**p**_0_, **p**_1_) ∈ M *for l* ∈ [0, 1]. *Here, we consider a Riemannian manifold of PD matrices*, 𝒮_++_. *Assume two PD matrices* **P**_0_, **P**_1_ ∈ 𝒮_++_. *Then, for l* ∈ [0, 1], *the geodesic curve joining* **P**_0_ *to* **P**_1_ *is defined as [69, Chapter. 6]:*

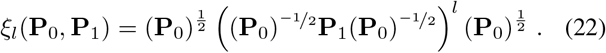

Note that **P**_0_ and **P**_1_ are obtained as the starting and end points of the geodesic path by choosing *l* = 0 and *l* = 1, respectively. The midpoint of the geodesic, obtained by setting 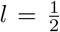, is called the *geometric mean*.

#### Definition 2

(Geodesic convexity). *Let* **p**_0_ *and* **p**_1_ *be two arbitrary points on a subset* 𝒜 *of a Riemannian manifold* ℳ. *Then a real-valued function f with domain* 𝒜 ⊂ ℳ *with f* : 𝒜 → ℝ *is called* geodesic convex *(g-convex) if the following relation holds:*

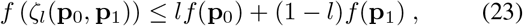

*where l* ∈ [0, 1] *and ζ*(**p**_0_, **p**_1_) *denotes the geodesic path connecting two points* **p**_0_ *and* **p**_1_ *as defined in Definition 1*.

The proof parallels the one provided in [70, Theorem. 3]:

*Proof*. First, we consider PD manifolds and express (23) in terms of geodesic paths and functions that lie on this particular space. We then show that 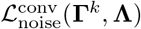 is strictly g-convex on this specific domain. Second, we then derive the update rule proposed in (12).

#### 1) G-convexity of the Majorizing Cost Function

Let *ξ*_*l*_(**Λ**_0_, **Λ**_1_) denote geodesics along the PD manifold as presented in Definition 1, and let define *f* (.) to be 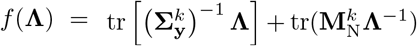, representing the cost function 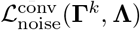.

We now show that *f* (**Λ**) is strictly g-convex on this specific domain. For continuous functions as considered in this paper, fulfilling (23) for *f* (**Λ**) and *ξ*_*l*_(**Λ**_0_, **Λ**_1_) with *l* = 1*/*2 is sufficient for strict g-convexity according to *mid-point convexity* [71]:

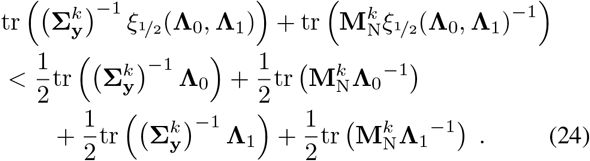

Given 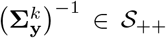, i.e., 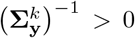 and the operator inequality [69, Chapter. 4]

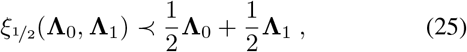

we have:

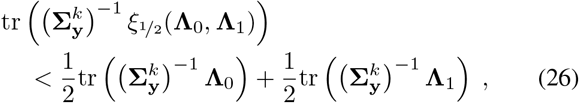

which is derived by multiplying both sides of (25) with 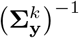 followed by taking the trace on both sides.

Similarly, we can write the operator inequality for 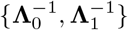 using (22) as:

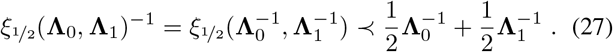

Multiplying both sides of (27) by 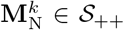 and applying the trace operator on both sides leads to:

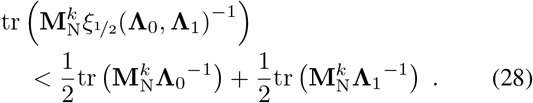

Summing up (26) and (28) proves inequality (24) and concludes the first part of the proof.

#### 2) Derivation of the Update Rule in (12)

We now present the second part of the proof by deriving the update rule in (12). Since the cost function 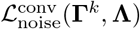 is strictly g-convex, its optimal solution in the *k*-th iteration is unique. More concretely, the optimum can be analytically derived by taking the derivative of (10) and setting the result to zero as follows:

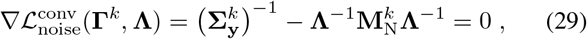

which results in

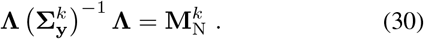

This solution is known as the *Riccati equation*, and is the geometric mean between 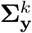 and 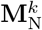 [72], [73]:

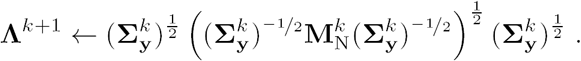

Deriving the update rule in (12) concludes the second part of the proof of Theorem 2.

### C. Proof of Theorem 3

We start the derivation of update rule (13) by constraining **Γ** to the set of diagonal matrices with non-negative entries 𝒮, i.e.,

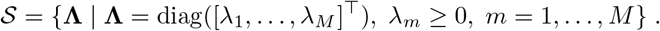

We continue by reformulating the constrained optimization with respect to the source covariance matrix,

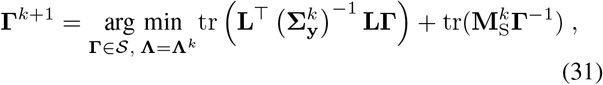

as follows:

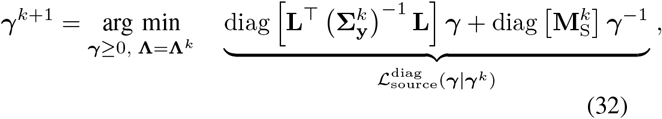

where 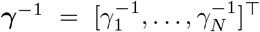 is defined as the elementwise inversion of ***γ***. Note that the set of diagonal matrices with all non-negative entries are positive semidefinite (PSD) by construction [74, Appendix A]. Thus, by constraining the space of solutions of optimization problem (31) to the set 𝒮, the PSD requirement for **Γ** reduces to the requirement that the diagonal elements of **Γ**, i.e., *γ*_*n*_, for *n* = 1, …, *N*, must be non-negative. The optimization with respect to the scalar source variances is then carried out by taking the derivative of (32) with respect to *γ*_*n*_, for *n* = 1, …, *N*, and setting it to zero, yields the following update rule:

**Γ**^*k*+1^ = diag(***γ***^*k*+1^), where,

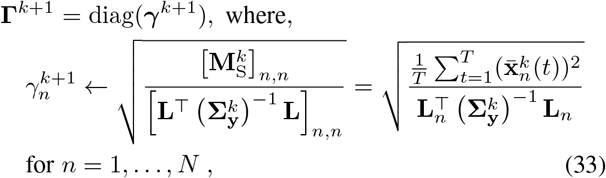

for *n* = 1, …, *N*, where **L**_*n*_ denotes the *n*-th column of the lead field matrix. Note that (33) is identical to the update rule of Champagne [28].

### D. Proof of Theorem 4

We prove Theorem 4 by showing that the alternating update rules for **Λ** and **Γ**, (12) and (13), are guaranteed to converge to a local minimum of the Bayesian Type-II likelihood (7). More generally, we prove that FUN learning is an instance of the general class of majorization-minimization (MM) algorithms, for which this property follows by construction. To this end, we first briefly review theoretical concepts behind the majorization-minimization (MM) algorithmic framework [62], [63] [61], [75].

#### 1) Required Conditions for Majorization-Minimization Algorithms

MM encompasses a family of iterative algorithms for optimizing general non-linear cost functions. The main idea behind MM is to replace the original cost function in each iteration by an upper bound, also known as majorizing function, whose minimum is easy to find. Interested readers are referred to [36] for an extensive list of applications on MM.

The problem of minimizing a continuous function *f* (**u**) within a closed convex set 𝒰 ⊂ ℝ^*n*^:

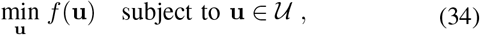

within the MM framework can be summarized as follows. First, construct a continuous *surrogate function g*(**u**|**u**^*k*^) that *majorizes*, or upper-bounds, the original function *f* (**u**) and coincides with *f* (**u**) at a given point **u**^*k*^:

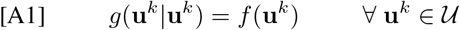

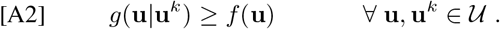

Second, starting from an initial value **u**^0^, generate a sequence of feasible points **u**^1^, **u**^2^, …, **u**^*k*^, **u**^*k*+1^ as solutions of a series of successive simple optimization problems, where

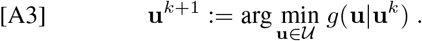

If a surrogate function fulfills conditions [A1]–[A3], then the value of the cost function *f* decreases in each iteration: *f* (**u**^*k*+1^) ≤ *f* (**u**^*k*^). For the smooth functions considered in this paper, we further require that the derivatives of the original and surrogate functions coincide at **u**^*k*^:

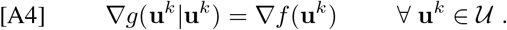

We can then formulate the following theorem:

##### Theorem 5.

*Assume that an MM algorithm fulfills conditions [A1]–[A4]. Then, every limit point of the sequence of minimizers generated in [A3], is a stationary point of the original optimization problem in* (34).

*Proof*. A detailed proof is provided in [63, Theorem 1].

*2) Details of the Proof of Theorem 4*

We now show that FUN learning is an instance of majorization-minimization as defined above, which fulfills Theorem 5.

*Proof*. We need to prove that conditions [A1]–[A4] are fulfilled for FUN learning. To this end, we recall the upper bound on log |**Σ**_**y**_| in (17), which fulfills condition [A2] since it majorizes log |**Σ**_**y**_| by virtue of the concavity of the log-determinant function and its first-order Taylor expansion around 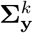. Besides, it automatically satisfies conditions [A1] and [A4] by construction, because the majorizing function in (17) is obtained through a Taylor expansion around 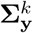. Concretely, [A1] is satisfied because the equality in (17) holds for 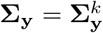. Similarly, [A4] is satisfied because the gradient of log |**Σ**_**y**_| at point 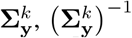 defines the linear Taylor approximation log 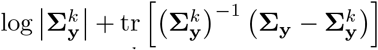. Thus, both gradients coincide in 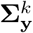 by construction. We can further prove that [A3] can be satisfied by showing that 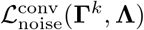 reaches its global minimum in each MM iteration. This is guaranteed if 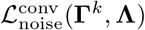 can be shown to be convex or g-convex with respect to **Λ**. To this end, we first require the subsequent proposition:

##### Proposition 1.

*Any local minimum of a g-convex function over a g-convex set is a global minimum*.

*Proof*. A detailed proof is presented in [76, Theorem 2.1].

Given Theorem 2, which already states that the costfunction 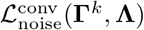 is g-convex, and Proposition 1, we can conclude that any local minimum of 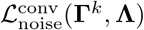 is a global minimum.

For brevity, we omit the proof of conditions [A1], [A2] and [A4] for the optimization with respect to **Γ** based on the convex surrogate function in (8), 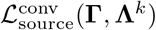, as it can be presented analogously. We here only show that [A3] is satisfied if 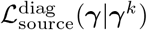 in (32) is a convex function with respect to ***γ***. Note that the g-convexity of 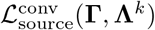 can also be proven using arguments analogous to those presented in appendix B.1. However, we instead prove a stronger condition, i.e., convexity, for simplifying the proof. To this end, we rewrite (32) as follows:

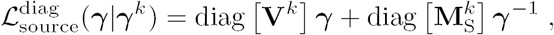

where 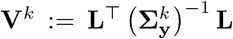 is defined as a parameter that does not depend on ***γ***. The convexity of 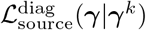 can be directly inferred from the convexity of diag [**V**^*k*^]***γ*** and diag 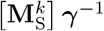 with respect to ***γ*** [77, Chapter. 3]. The convexity of 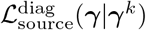, which ensures that condition [A3] can be satisfied using standard optimization, along with the fulfillment of conditions [A1], [A2] and [A4], ensure that

Theorem 5 holds for 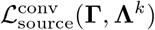. This completes the proof that the optimization of (7) with respect to **Γ** using the convex surrogate cost function (8) leads to an MM algorithm with convergence guarantees.

### E. Special Case of FUN Learning leads to Champagne with Heteroscedastic Noise Learning

We start by constraining **Λ** to the set of diagonal matrices with non-negative entries 𝒮, i.e.,

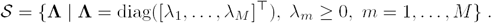

We then reformulate the constrained optimization with respect to the noise covariance matrix,

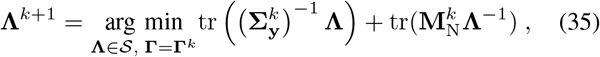

as follows:

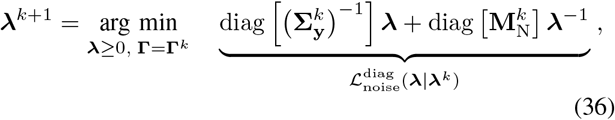

where 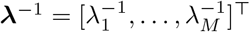 is defined as the element-wise inversion of ***λ***. Taking the derivative of (36) with respect to *λ*_*m*_, for *m* = 1, …, *M*, and setting it to zero, yields the following update rule:

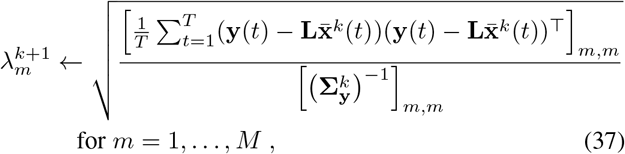

for *m* = 1, …, *M*, which is identical to the update rule of the Champagne with heteroscedastic noise learning as presented in [9].

## Acknowledgment

This result is part of a project that has received funding from the European Research Council (ERC) under the European Union’s Horizon 2020 research and innovation programme (Grant agreement No. 758985).

AH acknowledges scholarship support from the Machine Learning/Intelligent Data Analysis research group at Technische Universität Berlin. He further wishes to thank the Charité – Universitätsmedizin Berlin, the Berlin Mathematical School (BMS), and the Berlin Mathematics Research Center MATH+ for partial support. CC was supported by National Natural Science Foundation of China under Grants 62277023 and 62007013, as well as Hubei Provincial Natural Foundation of China under Grant 2021CFB384. KRM was partly funded by the German Ministry for Education and Research (under refs 01IS14013A-E, 01GQ1115, 01GQ0850, 01IS18056A, 01IS18025A and 01IS18037A), the German Research Foundation (DFG) as Math+: Berlin Mathematics Research Center (EXC 2046/1, project-ID: 390685689). Furthermore, KRM was partly supported by the Institute of Information & Communications Technology Planning & Evaluation (IITP) grants funded by the Korea Government (No. 2019-0-00079, Artificial Intelligence Graduate School Program, Korea University). SSN was funded in part by National Institutes of Health grants (R01DC004855, R01EB022717, R01DC176960, R01DC010145, R01NS100440, R01AG062196, and R01DC013979), University of California MRPI MRP-17–454755, the US Department of Defense grant (W81XWH-13-1-0494) and a research contract with Ricoh MEG USA Inc.

## References

[1] B. Rakitsch, C. Lippert, K. Borgwardt, and O. Stegle, “It is all in the noise: Efficient multi-task gaussian process inference with structured residuals,” in Proceedings of the 26th International Conference on Neural Information Processing Systems-Volume 1, 2013, pp. 1466–1474.

[2] M. Cai, N. W. Schuck, J. W. Pillow, and Y. Niv, “A Bayesian method for reducing bias in neural representational similarity analysis,” in Advances in Neural Information Processing Systems, 2016, pp. 4951–4959.

[3] M. B. Cai, M. Shvartsman, A. Wu, H. Zhang, and X. Zhu, “Incorporating structured assumptions with probabilistic graphical models in fMRI data analysis,” Neuropsychologia, p. 107500, 2020.

[4] S. Liu, J. Jia, Y. D. Zhang, and Y. Yang, “Image reconstruction in electrical impedance tomography based on structure-aware sparse Bayesian learning,” IEEE Transactions on Medical Imaging, vol. 37, no. 9, pp. 2090–2102, 2018.

[5] S. Liu, Y. Huang, H. Wu, C. Tan, and J. Jia, “Efficient multitask structure-aware sparse Bayesian learning for frequency-difference electrical impedance tomography,” IEEE Transactions on Industrial Informatics, vol. 17, no. 1, pp. 463–472, 2020.

[6] S. Hannan, M. Faulkner, K. Aristovich, J. Avery, M. C. Walker, and D. S. Holder, “In vivo imaging of deep neural activity from the cortical surface during hippocampal epileptiform events in the rat brain using electrical impedance tomography,” NeuroImage, vol. 209, p. 116525, 2020.

[7] Q. Bertrand, M. Massias, A. Gramfort, and J. Salmon, “Handling correlated and repeated measurements with the smoothed multivariate square-root Lasso,” in Advances in Neural Information Processing Systems, 2019, pp. 3959–3970.

[8] A. Hashemi, C. Cai, G. Kutyniok, K.-R. Müller, S. Nagarajan, and S. Haufe, “Unification of sparse Bayesian learning algorithms for electromagnetic brain imaging with the majorization minimization framework,” NeuroImage, vol. 239, p. 118309, 2021.

[9] C. Cai, A. Hashemi, M. Diwakar, S. Haufe, K. Sekihara, and S. S. Nagarajan, “Robust estimation of noise for electromagnetic brain imaging with the Champagne algorithm,” NeuroImage, vol. 225, p. 117411, 2021.

[10] D. P. Wipf and B. D. Rao, “An empirical Bayesian strategy for solving the simultaneous sparse approximation problem,” IEEE Transactions on Signal Processing, vol. 55, no. 7, pp. 3704–3716, 2007.

[11] Z. Zhang and B. D. Rao, “Sparse signal recovery with temporally correlated source vectors using sparse Bayesian learning,” IEEE Journal of Selected Topics in Signal Processing, vol. 5, no. 5, pp. 912–926, 2011.

[12] S. Van de Geer, J. Lederer et al., “The Lasso, correlated design, and improved oracle inequalities,” in From Probability to Statistics and Back: High-Dimensional Models and Processes–A Festschrift in Honor of Jon A. Wellner. Institute of Mathematical Statistics, 2013, pp. 303–316.

[13] A. Dalalyan, M. Hebiri, K. Meziani, and J. Salmon, “Learning heteroscedastic models by convex programming under group sparsity,” in International Conference on Machine Learning, 2013, pp. 379–387.

[14] J. Lederer and C. L. Muller, “Don’t fall for tuning parameters: tuning-free variable selection in high dimensions with the TREX,” in Proceedings of the Twenty-Ninth AAAI Conference on Artificial Intelligence, 2015, pp. 2729–2735.

[15] E. J. Candés, J. K. Romberg, and T. Tao, “Stable signal recovery from incomplete and inaccurate measurements,” Communications on pure and applied mathematics, vol. 59, no. 8, pp. 1207–1223, 2006.

[16] D. L. Donoho, “Compressed sensing,” IEEE Transactions on Information Theory, vol. 52, no. 4, pp. 1289–1306, 2006.

[17] D. Malioutov, M. Cetin, and A. S. Willsky, “A sparse signal reconstruction perspective for source localization with sensor arrays,” IEEE Transactions on Signal Processing, vol. 53, no. 8, pp. 3010–3022, 2005.

[18] T. Li and A. Nehorai, “Maximum likelihood direction finding in spatially colored noise fields using sparse sensor arrays,” IEEE Transactions on Signal Processing, vol. 59, no. 3, pp. 1048–1062, 2010.

[19] C. E. Chen, F. Lorenzelli, R. E. Hudson, and K. Yao, “Stochastic maximum-likelihood DOA estimation in the presence of unknown nonuniform noise,” IEEE Transactions on Signal Processing, vol. 56, no. 7, pp. 3038–3044, 2008.

[20] M. S. Zhdanov, Inverse theory and applications in geophysics. Elsevier, 2015, vol. 36.

[21] S. F. Cotter, B. D. Rao, K. Engan, and K. Kreutz-Delgado, “Sparse solutions to linear inverse problems with multiple measurement vectors,” IEEE Transactions on Signal Processing, vol. 53, no. 7, pp. 2477–2488, 2005.

[22] M. Hämäläinen, R. Hari, R. J. Ilmoniemi, J. Knuutila, and O. V. Lounasmaa, “Magnetoencephalography—theory, instrumentation, and applications to noninvasive studies of the working human brain,” Reviews of modern Physics, vol. 65, no. 2, p. 413, 1993.

[23] R. D. Pascual-Marqui, C. M. Michel, and D. Lehmann, “Low resolution electromagnetic tomography: a new method for localizing electrical activity in the brain,” International Journal of psychophysiology, vol. 18, no. 1, pp. 49–65, 1994.

[24] I. F. Gorodnitsky, J. S. George, and B. D. Rao, “Neuromagnetic source imaging with FOCUSS: a recursive weighted minimum norm algorithm,” Electroencephalography and Clinical Neurophysiology, vol. 95, no. 4, pp. 231–251, 1995.

[25] S. Haufe, V. V. Nikulin, A. Ziehe, K.-R. Müller, and G. Nolte, “Combining sparsity and rotational invariance in EEG/MEG source reconstruction,” NeuroImage, vol. 42, no. 2, pp. 726–738, 2008.

[26] A. Gramfort, M. Kowalski, and M. Hämäläinen, “Mixed-norm estimates for the M/EEG inverse problem using accelerated gradient methods,” Physics in Medicine and Biology, vol. 57, no. 7, p. 1937, 2012.

[27] S. Castaño-Candamil, J. Höhne, J.-D. Martínez-Vargas, X.-W. An, G. Castellanos-Domínguez, and S. Haufe, “Solving the EEG inverse problem based on space–time–frequency structured sparsity constraints,” NeuroImage, vol. 118, pp. 598–612, 2015.

[28] D. Wipf and S. Nagarajan, “A unified Bayesian framework for MEG/EEG source imaging,” NeuroImage, vol. 44, no. 3, pp. 947–966, 2009.

[29] M. W. Seeger and D. P. Wipf, “Variational Bayesian inference techniques,” IEEE Signal Processing Magazine, vol. 27, no. 6, pp. 81–91, 2010.

[30] W. Wu, S. Nagarajan, and Z. Chen, “Bayesian machine learning: EEG\MEG signal processing measurements,” IEEE Signal Processing Magazine, vol. 33, no. 1, pp. 14–36, 2016.

[31] D. P. Wipf and B. D. Rao, “Sparse Bayesian learning for basis selection,” IEEE Transactions on Signal Processing, vol. 52, no. 8, pp. 2153–2164, 2004.

[32] M. E. Tipping, “Sparse Bayesian learning and the relevance vector machine,” Journal of Machine Learning Research, vol. 1, no. Jun, pp. 211–244, 2001.

[33] S. Mika, G. Rätsch, and K.-R. Müller, “A mathematical programming approach to the kernel fisher algorithm,” Advances in Neural Information Processing Systems, vol. 13, pp. 591–597, 2001.

[34] K. Sekihara and S. S. Nagarajan, Electromagnetic brain imaging: a Bayesian perspective. Springer, 2015.

[35] D. P. Wipf, J. P. Owen, H. T. Attias, K. Sekihara, and S. S. Nagarajan, “Robust Bayesian estimation of the location, orientation, and time course of multiple correlated neural sources using MEG,” NeuroImage, vol. 49, no. 1, pp. 641–655, 2010.

[36] Y. Sun, P. Babu, and D. P. Palomar, “Majorization-minimization algorithms in signal processing, communications, and machine learning,” IEEE Transactions on Signal Processing, vol. 65, no. 3, pp. 794–816, 2017.

[37] P. Petersen, S. Axler, and K. Ribet, Riemannian geometry. Springer, 2006, vol. 171.

[38] S. Haufe and A. Ewald, “A simulation framework for benchmarking EEG-based brain connectivity estimation methodologies,” Brain topography, pp. 1–18, 2016.

[39] Y. Huang, L. C. Parra, and S. Haufe, “The New York head — a precise standardized volume conductor model for EEG source localization and tES targeting,” NeuroImage, vol. 140, pp. 150–162, 2016.

[40] Y. Rubner, C. Tomasi, and L. J. Guibas, “The earth mover’s distance as a metric for image retrieval,” International Journal of Computer Vision, vol. 40, no. 2, pp. 99–121, 2000.

[41] N. Chinchor and B. M. Sundheim, “Muc-5 evaluation metrics,” in Fifth Message Understanding Conference (MUC-5), 1993.

[42] S. S. Dalal, J. Zumer, V. Agrawal, K. Hild, K. Sekihara, and S. Nagarajan, “NUTMEG: a neuromagnetic source reconstruction toolbox,” Neurology & Clinical Neurophysiology: NCN, vol. 2004, p. 52, 2004.

[43] L. B. Hinkley, C. L. Dale, T. L. Luks, A. M. Findlay, P. Bukshpun, N. Pojman, T. Thieu, W. K. Chung, J. Berman, T. P. Roberts et al., “Sensorimotor cortical oscillations during movement preparation in 16p11. 2 deletion carriers,” Journal of Neuroscience, vol. 39, no. 37, pp. 7321–7331, 2019.

[44] L. B. Hinkley, E. De Witte, M. Cahill-Thompson, D. Mizuiri, C. Garrett, S. Honma, A. Findlay, M. L. Gorno-Tempini, P. Tarapore, H. E. Kirsch et al., “Optimizing magnetoencephalographic imaging estimation of language lateralization for simpler language tasks,” Frontiers in Human Neuroscience, vol. 14, p. 105, 2020.

[45] S. S. Dalal, J. M. Zumer, A. G. Guggisberg, M. Trumpis, D. D. Wong, K. Sekihara, and S. S. Nagarajan, “MEG/EEG source reconstruction, statistical evaluation, and visualization with NUTMEG,” Computational Intelligence and Neuroscience, vol. 2011, 2011.

[46] J. P. Owen, D. P. Wipf, H. T. Attias, K. Sekihara, and S. S. Nagarajan, “Performance evaluation of the Champagne source reconstruction algorithm on simulated and real M/EEG data,” Neuroimage, vol. 60, no. 1, pp. 305–323, 2012.

[47] C. Cai, M. Diwakar, D. Chen, K. Sekihara, and S. S. Nagarajan, “Robust empirical Bayesian reconstruction of distributed sources for electromagnetic brain imaging,” IEEE Transactions on Medical Imaging, vol. 39, no. 3, pp. 567–577, 2019.

[48] K. Matsuura and Y. Okabe, “Selective minimum-norm solution of the biomagnetic inverse problem,” IEEE Transactions on Biomedical Engineering, vol. 42, no. 6, pp. 608–615, 1995.

[49] R. D. Pascual-Marqui, “Discrete, 3D distributed, linear imaging methods of electric neuronal activity. Part 1: exact, zero error localization,” 2007.

[50] R. D. Pascual-Marqui et al., “Standardized low-resolution brain electromagnetic tomography (sLORETA): technical details,” Methods Find Exp Clin Pharmacol, vol. 24, no. Suppl D, pp. 5–12, 2002.

[51] B. D. Van Veen, W. Van Drongelen, M. Yuchtman, and A. Suzuki, “Localization of brain electrical activity via linearly constrained minimum variance spatial filtering,” IEEE Transactions on Biomedical Engineering, vol. 44, no. 9, pp. 867–880, 1997.

[52] D. A. Engemann and A. Gramfort, “Automated model selection in covariance estimation and spatial whitening of MEG and EEG signals,” NeuroImage, vol. 108, pp. 328–342, 2015.

[53] H. M. Huizenga, J. C. De Munck, L. J. Waldorp, and R. P. Grasman, “Spatiotemporal EEG/MEG source analysis based on a parametric noise covariance model,” IEEE Transactions on Biomedical Engineering, vol. 49, no. 6, pp. 533–539, 2002.

[54] J. C. De Munck, H. M. Huizenga, L. J. Waldorp, and R. Heethaar, “Estimating stationary dipoles from MEG/EEG data contaminated with spatially and temporally correlated background noise,” IEEE Transactions on Signal Processing, vol. 50, no. 7, pp. 1565–1572, 2002.

[55] F. Bijma, J. C. De Munck, H. M. Huizenga, and R. M. Heethaar, “A mathematical approach to the temporal stationarity of background noise in MEG/EEG measurements,” NeuroImage, vol. 20, no. 1, pp. 233–243, 2003.

[56] J. C. De Munck, F. Bijma, P. Gaura, C. A. Sieluzycki, M. I. Branco, and R. M. Heethaar, “A maximum-likelihood estimator for trial-to-trial variations in noisy MEG/EEG data sets,” IEEE Transactions on Biomedical Engineering, vol. 51, no. 12, pp. 2123–2128, 2004.

[57] S. C. Jun, S. M. Plis, D. M. Ranken, and D. M. Schmidt, “Spatiotemporal noise covariance estimation from limited empirical magnetoencephalo-graphic data,” Physics in Medicine & Biology, vol. 51, no. 21, p. 5549, 2006.

[58] M. Massias, O. Fercoq, A. Gramfort, and J. Salmon, “Generalized concomitant multi-task lasso for sparse multimodal regression,” in International Conference on Artificial Intelligence and Statistics, 2018, pp. 998–1007.

[59] C. Cai, L. Hinkley, Y. Gao, A. Hashemi, S. Haufe, K. Sekihara, and S. S. Nagarajan, “Empirical bayesian localization of event-related time-frequency neural activity dynamics,” NeuroImage, vol. 258, p. 119369, 2022.

[60] M. Shvartsman, N. Sundaram, M. Aoi, A. Charles, T. Willke, and J. Cohen, “Matrix-normal models for fMRI analysis,” in International Conference on Artificial Intelligence and Statistics. PMLR, 2018, pp. 1914–1923.

[61] T. T. Wu, K. Lange et al., “The MM alternative to EM,” Statistical Science, vol. 25, no. 4, pp. 492–505, 2010.

[62] M. W. Jacobson and J. A. Fessler, “An expanded theoretical treatment of iteration-dependent majorize-minimize algorithms,” IEEE Transactions on Image Processing, vol. 16, no. 10, pp. 2411–2422, 2007.

[63] M. Razaviyayn, M. Hong, and Z.-Q. Luo, “A unified convergence analysis of block successive minimization methods for nonsmooth optimization,” SIAM Journal on Optimization, vol. 23, no. 2, pp. 1126–1153, 2013.

[64] A. Hashemi, C. Cai, Y. Gao, S. Ghosh, K.-R. Müller, S. S. Nagarajan, and S. Haufe, “Joint learning of full-structure noise in hierarchical Bayesian regression models,” bioRxiv, 2022.

[65] A. Breloy, Y. Sun, P. Babu, G. Ginolhac, and D. P. Palomar, “Robust rank constrained kronecker covariance matrix estimation,” in 2016 50th Asilomar Conference on Signals, Systems and Computers. IEEE, 2016, pp. 810–814.

[66] B. Xin, Y. Wang, W. Gao, and D. Wipf, “Building invariances into sparse subspace clustering,” IEEE Transactions on Signal Processing, vol. 66, no. 2, pp. 449–462, 2017.

[67] P. Babu, “MELT—maximum-likelihood estimation of low-rank Toeplitz covariance matrix,” IEEE Signal Processing Letters, vol. 23, no. 11, pp. 1587–1591, 2016.

[68] A. Hashemi, Y. Gao, C. Cai, S. Ghosh, K. R. Müller, S. S. Nagarajan, and S. Haufe, “Efficient hierarchical Bayesian inference for spatio-temporal regression models in neuroimaging,” in Thirty-Fifth Conference on Neural Information Processing Systems, 2021.

[69] R. Bhatia, Positive definite matrices. Princeton University Press, 2009, vol. 24.

[70] P. Zadeh, R. Hosseini, and S. Sra, “Geometric mean metric learning,” in International Conference on Machine Learning, 2016, pp. 2464–2471.

[71] C. Niculescu and L.-E. Persson, Convex functions and their applications. Springer, 2006.

[72] J. V. Davis, B. Kulis, P. Jain, S. Sra, and I. S. Dhillon, “Information-theoretic metric learning,” in Proceedings of the 24th International Conference on Machine Learning, 2007, pp. 209–216.

[73] S. Bonnabel and R. Sepulchre, “Riemannian metric and geometric mean for positive semidefinite matrices of fixed rank,” SIAM Journal on Matrix Analysis and Applications, vol. 31, no. 3, pp. 1055–1070, 2009.

[74] E. De Klerk, Aspects of semidefinite programming: interior point algorithms and selected applications. Springer Science & Business Media, 2006, vol. 65.

[75] D. R. Hunter and K. Lange, “A tutorial on MM algorithms,” The American Statistician, vol. 58, no. 1, pp. 30–37, 2004.

[76] T. Rapcsak, “Geodesic convexity in nonlinear optimization,” Journal of Optimization Theory and Applications, vol. 69, no. 1, pp. 169–183, 1991.

[77] S. P. Boyd and L. Vandenberghe, Convex optimization. Cambridge university press, 2004.

